# Aβ toxicity rescued by protein retention in the ER

**DOI:** 10.1101/2021.08.18.456775

**Authors:** James H Catterson, Lucy Minkley, Salomé Aspe, Sebastian Judd-Mole, Sofia Moura, Miranda C Dyson, Arjunan Rajasingam, Nathaniel S Woodling, Magda L Atilano, Mumtaz Ahmad, Claire S Durrant, Tara L Spires-Jones, Linda Partridge

## Abstract

Accumulation of Aβ in the brain is one of the hallmarks of Alzheimer’s disease (AD). In the adult *Drosophila* brain, human Aβ over-expression is toxic and leads to deterioration of climbing ability and shortened lifespan. However, it remains unknown if Aβ is inherently toxic or if it triggers toxic downstream pathways that lead to neurodegeneration. Here, we describe a novel, and previously unidentified, protective role of intracellular laminin chain accumulation. Despite high Aβ levels, over-expression of the extracellular matrix protein subunit Laminin B1 (LanB1) resulted in a robust rescue of toxicity, highlighting a potential protective mechanism of resistance to Aβ. Over-expression of other Laminin subunits and a Collagen IV subunit also significantly rescued Aβ toxicity, while combining LanB1 with these subunits led to an even larger rescue. Imaging revealed that LanB1 was retained in the ER but had no effect on the secretion of Aβ into the extracellular milieu. LanB1 rescued toxicity independently of the IRE1α/XBP1-mediated branch of the ER stress response. Interestingly, over-expression of ER-targeted GFP also rescued Aβ toxicity, indicating a potentially broader benefit of ER protein retention. Finally, in proof-of-principle lentiviral transduction experiments using murine organotypic hippocampal slice cultures, over-expression of mouse Lamb1 resulted in ER-retention in transduced cells, highlighting a conserved mechanism. Typically, retention of proteins in the ER is detrimental to cellular health, but in the context of neuronal Aβ toxicity it may prove to be beneficial and a new therapeutic avenue for AD.

## Introduction

Despite decades of intensive research and a large number of clinical trials, Alzheimer’s disease (AD) has no cure. Most cases are sporadic, with age being the biggest risk factor. The neuropathological hallmarks include accumulation of Amyloid β (Aβ) plaques and neurofibrillary tau tangles^1^. AD is thought to be triggered by the accumulation of Aβ peptides, derived from the misprocessing of amyloid precursor protein (APP), resulting in increased cellular stress, accumulation of toxic hyperphosphorylated tau, and eventual neuronal cell death^2^. Characteristic features of AD include aggregates of unfolded proteins, increased reactive oxygen species and metabolic dysregulation in the affected neurons^3,4^. The endoplasmic reticulum (ER) recognizes these alterations in neuronal homeostasis, and consequently AD brains display many signs of ER stress^5,6^, which appears early in AD progression^7,8^. ER stress generally occurs upon the accumulation of unfolded or misfolded proteins in the ER lumen. The ER chaperone BiP (Grp78) dissociates from membrane sensors (Inositol-requiring enzyme 1α (IRE1α), Protein kinase-like endoplasmic reticulum kinase (PERK), and Activating Transcription Factor 6 (ATF6)), and binds to these misfolded proteins^9^. With the removal of BiP, IRE1α oligomerises then catalyses the “unconventional” splicing of X-box binding protein 1 (Xbp1) mRNA, generating XBP1s, a transcription factor that regulates genes involved in the ER stress response, including BiP^10^.

Aβ pathology is largely extracellular but there is also a lively debate^11,12^ in the AD field about intracellular Aβ, which may play a major role in neurodegeneration^13,14^. Although the existence of intracellular Aβ has become less controversial, its role in the induction of the ER stress response is unresolved, and the mechanisms by which protein misfolding contributes to AD pathogenesis are still unclear^15^.

The detailed molecular mechanisms underlying the aetiology of AD remain to be confirmed, and many *in vitro* and *in vivo* models have been developed to study them^16,17^. *Drosophila melanogaster* has been widely used as a model system to study neurodegenerative disorders, including AD^16^. In the adult fly brain, neuronally expressed Aβ is predominantly found in the cell body of neurons, but can also be observed in the neuropil when glial function is affected^18^, indicating that glia, which account for ∼10% of the cells^19^ yet cover a large area of the adult fly brain^20^, play an important role clearing Aβ.

In flies, the ER stress response is activated in response to neuronal Aβ, with increased Xbp1 splicing and BiP levels^21–23^. Indeed, Xbp1 acts to buffer the toxic effects of Aβ since over-expression of spliced Xbp1 reduces Aβ levels and partially rescues toxicity, while Xbp1 knockdown increases Aβ levels and exacerbates toxicity^21,23^.

The ER is involved in the secretion of many proteins, including those targeted to the extracellular milieu, such as collagens and laminins^24^. The assembly of laminins and collagens from their individual subunits occurs in the ER before secretion into the trans-Golgi network and then toward their final destination outside the cell, where they form the extracellular matrix (ECM)^25^. Laminins are obligate heterotrimers consisting of large α, β, and γ subunits that combine via the triple-helical coiled-coil domain in the centre of each chain to form cruciform-shaped structures. The expression of the β1 subunit of the ECM protein Laminin (LamB1) appears restricted to regions of the brain that are susceptible to neurodegeneration, especially the hippocampal tri-synaptic circuit^26,27^. Indeed, modulating LamB1 expression has been associated with changes in memory formation^28^, while Aβ-induced memory deficits are rescued by LamB1 modulation^29^. Under normal conditions laminin interacts with APP^30^. Additionally, anti-laminin immunoreactivity levels in human cerebrospinal fluid have been shown to correlate with the pathogenesis of AD and vascular dementia^31^. *In vitro,* the laminin heterotrimer can inhibit Aβ fibrillation and depolymerise Aβ, although without reduction of toxicity^32–34^. Notably, Aβ can induce memory deficits that can be rescued by ECM manipulation^29,35–37^. The complexity of neuron-matrix interactions makes it difficult to recapitulate ECM organisation and function in cell culture. It is, therefore, important to design experiments to evaluate these interactions *in vivo*.

Laminin in the brain is typically found extracellularly at basement membranes in the vasculature, and at the blood brain barrier^38^. However, synaptic laminin has also been characterised at the neuromuscular junction (NMJ)^39,40^. In the hippocampus, synaptic α5-laminin was found to be necessary and sufficient to stabilize dendritic spine dynamics^41^. Surprisingly, intraneuronal laminin expression has also been identified in the hippocampus and may have functions other than extracellular structural integrity^28,42^. Indeed, laminin has been observed in hippocampal neuronal perikarya and other susceptible regions of the brain important in the development of AD^43,44^. Laminin expression was also observed to be increased in the AD prefrontal cortex compared to non-disease controls^45^. Similarly, Collagen VI α1 chain was upregulated in dentate gyrus soma in mice expressing hAPP, and could rescue Aβ toxicity *in vitro* by sequestering Aβ into large aggregates in the extracellular milieu^35^.

Vertebrates have multiple genes for all three laminin subunits; five α-chains (α1–α5), four β-chains (β1–β4) and three γ-chains (γ1–γ3) are known, and they combine to form at least 16 different laminin heterotrimers^46^, not including novel spliced forms^47^. In *Drosophila*, there are two α (LanA and wb), one β (LanB1), and one γ (LanB2) subunits, resulting in only two laminin heterotrimers^48^ (diagram in **Fig. 1h**). Laminin trimerization occurs in the ER where the β and γ subunits assemble and require α subunit incorporation to form a functional laminin heterotrimer before secretion^49–51^. If either the β or γ subunit is missing, the other accumulates intracellularly and the α subunit can be secreted as a monomer. Knockdown of α laminin also results in β and γ subunit retention in the ER^52^. Similarly, over-expression of laminin monomers in glial cells leads to ER expansion and triggers ER stress, impairing correct development and locomotion in *Drosophila* larvae^53^. Collagen IV is another obligate heterotrimer that can accumulate intracellularly when subunit stoichiometry is altered^54^.

**Fig. 1.**
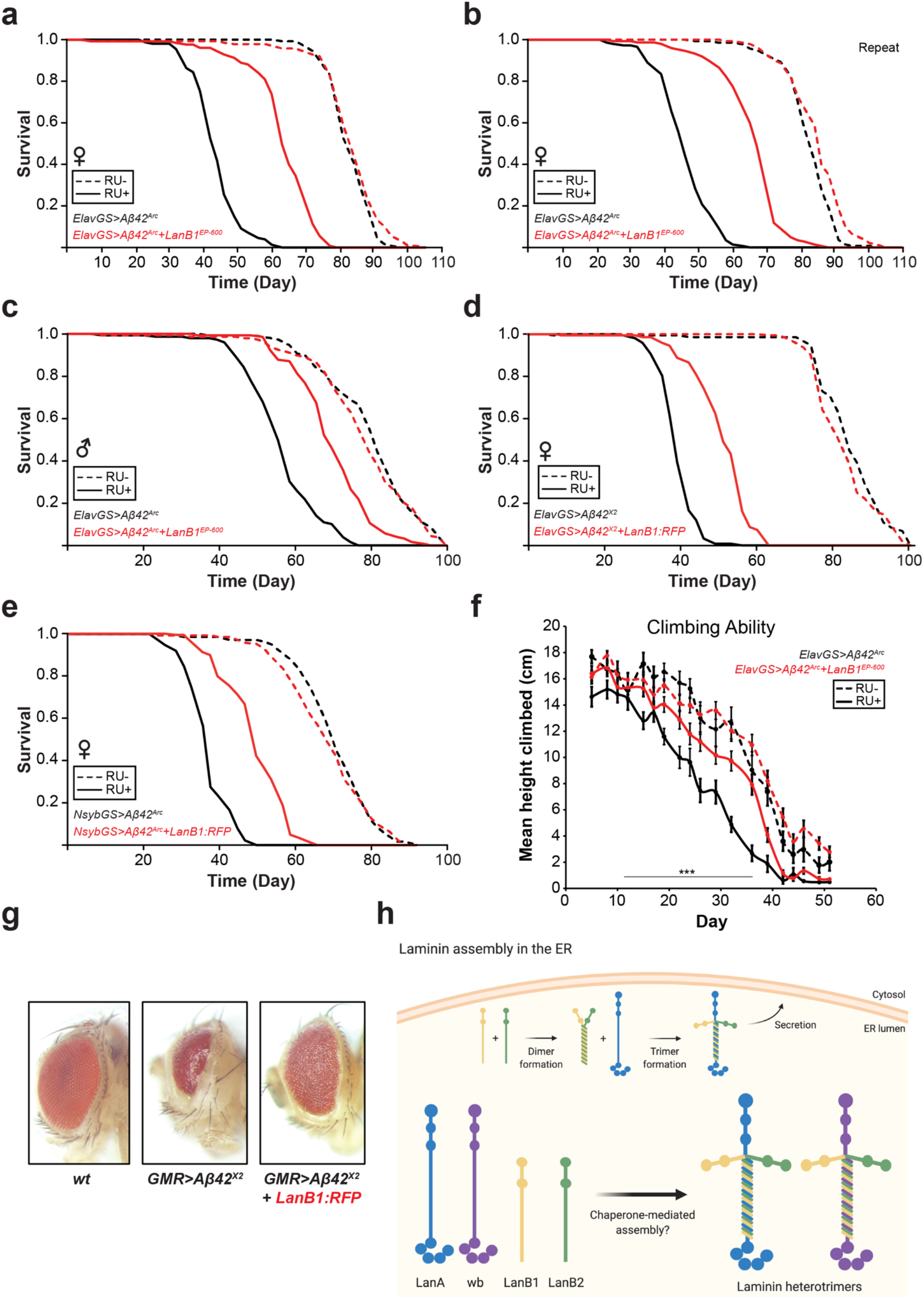
Co-expression of LanB1 rescued the toxic effect of Aβ expression. **a** Survival curves of female flies expressing Aβ^Arc^ in adult neurons. Induction of Aβ^Arc^ with ElavGS significantly (p = 1.31 x 10^−70^; log rank test) shortened lifespan compared to uninduced controls. LanB1 and Aβ^Arc^ co-expression resulted in a significant rescue (p = 1.38 x 10^−50^; log rank test) of the short-lived phenotype. **b** Repeat of experiment in **a**. LanB1 significantly rescued Aβ^Arc^ toxicity (p = 8.31 x 10^−56^; log rank test). **c** LanB1 significantly rescued Aβ^Arc^ toxicity in male flies (p = 8.31 x 10^−56^; log rank test). **d** Induction of Aβ^X2^ significantly (p = 5.99 x 10^−69^; log rank test) shortened lifespan compared to uninduced controls. LanB1 and Aβ^X2^ co-expression resulted in a significant rescue (p = 8.04 x 10^−40^; log rank test). **e** Induction of Aβ^Arc^ with NsybGS significantly (p = 3.38 x 10^−67^; log rank test) shortened lifespan compared to uninduced controls. LanB1 significantly rescued Aβ^Arc^ toxicity (p = 6.03 x 10^−40^; log rank test). Dashed lines represent uninduced ‘RU-‘ controls, solid lines represent induced ‘RU+’ conditions. For all lifespan experiments n = 150 flies per condition. **f** Climbing ability was measured until day 51. Climbing ability of Aβ^Arc^ flies was lower than that of all other groups (p < 0.0001, two-way ANOVA with Tukey’s post-hoc test). LanB1 significantly rescued the Aβ^Arc^-induced decline in climbing ability, though not to the same level as uninduced controls (p < 0.0001, two-way ANOVA with Tukey’s post-hoc test). Data are shown as mean ± SEM (n = 37-70 flies measured per time point). **g** LanB1 suppressed Aβ toxicity in the eyes of flies raised at 29°C. Control flies displayed a highly ordered ommatidia lattice (left). Expression of Aβ^X2^ using the eye-specific GMR-GAL4 driver resulted in small, glassy eyes that accumulated necrotic spots (middle). Co-expression of Aβ^X2^ with LanB1 rescued most of the size and organisation defects (right). **h** *Drosophila* laminins and their assembly in the ER. Created using BioRender.

If ER stress is part of the AD pathogenic cascade, and it is also activated in response to modulation of laminin/collagen subunit stoichiometry, then experimentally increasing ectopic ECM protein subunit expression in adult neurons might be expected to overwhelm the ER and exacerbate AD pathogenesis. On the contrary, we found that neuronal over-expression of *Drosophila* laminin and collagen subunits led to pronounced intra-ER accumulation of these subunits, yet could robustly ameliorate the toxic effects of Aβ, such as reduced lifespan and climbing ability, without reducing Aβ levels. We also found that over-expression of mouse Lamb1 in murine organotypic hippocampal slice cultures resulted in ER-retention of these monomers, indicating a conserved mechanism. Interestingly, over-expression of ER-targeted GFP also rescued Aβ toxicity, indicating a potentially broader benefit of ER protein retention. Protein accumulation in the ER is typically seen as detrimental to cellular health, but in the context of neuronal Aβ toxicity it may prove to be beneficial.

## Results

### Over-expression of the laminin β-chain LanB1 rescues the toxic effect of neuronal Aβ42 expression

To test if modulation of ECM components could alter the toxicity of Aβ_42_ *in vivo*, we took advantage of the Aβ_42_-shortened lifespan of *Drosophila* as a read-out. We used a drug-inducible *Drosophila* AD model that expressed the highly aggregative Arctic Aβ_42_ (Aβ^Arc^)^55^, which when expressed in the neurons of adult flies shortens lifespan and induces behavioural defects and neurodegeneration^21,56^. The Aβ sequence contains a signal peptide sequence from the *necrotic* gene^55^ to target it to the secretory pathway, resulting in Aβ that is secreted into the extracellular milieu. The pan-neuronal Elav-GeneSwitch (ElavGS) driver can be induced by RU486 (‘RU’, a steroid drug inducer) to switch on gene expression. We generated ElavGS; Aβ^Arc^ recombinant flies and used these to screen ECM-related genes by driving co-expression in neurons during adulthood. We observed a pronounced rescue of lifespan with co-expression of Aβ and the β subunit of Laminin (LanB1), using the EP-600 line with a P-element inserted 548 bp upstream of the *LanB1* gene^48^, compared to the Aβ^Arc^-alone controls (**Supplementary Fig. 1a**). We examined further Laminin transgenic lines and found that two independent LanB2 RNAi lines also significantly ameliorated Aβ^Arc^ toxicity, but not to the same extent as LanB1 co-expression (**Supplementary Fig. 1b**). As LanB1 co-expression induced the most consistent and pronounced rescue of Aβ^Arc^ toxicity, we took this line forward for further study.

Previous studies have shown that during *Drosophila* development laminin is predominantly produced by the fat body, haemocytes, and glia^53,57,58^. Information about laminin expression in the adult fly brain is less abundant. However, by examining gene expression in the adult fly brain using *SCope*^59^, a single-cell transcriptome atlas of the adult *Drosophila* brain, we found that expression of laminin subunits appeared to be specific to haemocytes and a subset of glial cells, while endogenous expression in neurons was low (**Supplementary Fig. 2a** and **b**), indicating that neuronal expression of these subunits is mainly ectopic.

To establish the robustness of the rescue of Aβ toxicity and to eliminate potential confounding effects of genetic background, we tested further fly lines in the standardised Dahomey (*w^Dah^*) genetic background. The UAS-LanB1^EP-600^ mutant was backcrossed, and the lifespan rescue experiment repeated, further including the genetically identical, uninduced controls. Co-expression of LanB1 and Aβ^Arc^ in females resulted in a significant rescue of the short-lived phenotype (**Fig. 1a**, repeat experiments, **Fig. 1b** and **Supplementary Fig. 1c**). The rescue was also significant in male flies (**Fig. 1c**). We next assessed the ability of LanB1 co-expression to rescue the shortened lifespan in a different fly AD model, in which two copies of wild-type, tandem Aβ_42_ (Aβ^X2^)^21^ with an export sequence from the *argos* gene, are expressed in adult neurons. Co-expression of an RFP-tagged LanB1 significantly rescued the short lifespan with Aβ^X2^ induction (**Fig. 1d** and **Supplementary Fig. 1d**). To assess if these neuronal effects were specific to the ElavGS driver, we used a second, independent, pan-neuronal GeneSwitch driver, NsybGS. LanB1 significantly rescued the shortened lifespan with Aβ^Arc^ induction in NsybGS flies (**Fig. 1e**).

To assess the effects of LanB1 on neuromuscular performance in AD flies, we measured their climbing behaviour. Expression of Aβ^Arc^ in adult neurons led to a significant decline in climbing ability compared to uninduced controls (**Fig. 1f**), and LanB1 co-expression significantly rescued the deficit. We also examined the feeding behaviour of the flies, using the Capillary Feeder (CAFE) assay^60^. Flies were induced/uninduced for 3 weeks and then feeding behaviour was measured with all flies eating food without RU. Aβ^Arc^ flies ingested a significantly smaller amount of food over 7 days compared to uninduced controls, while LanB1 co-expression completely rescued feeding behaviour to uninduced levels (**Fig. 2e**).

**Fig. 2.**
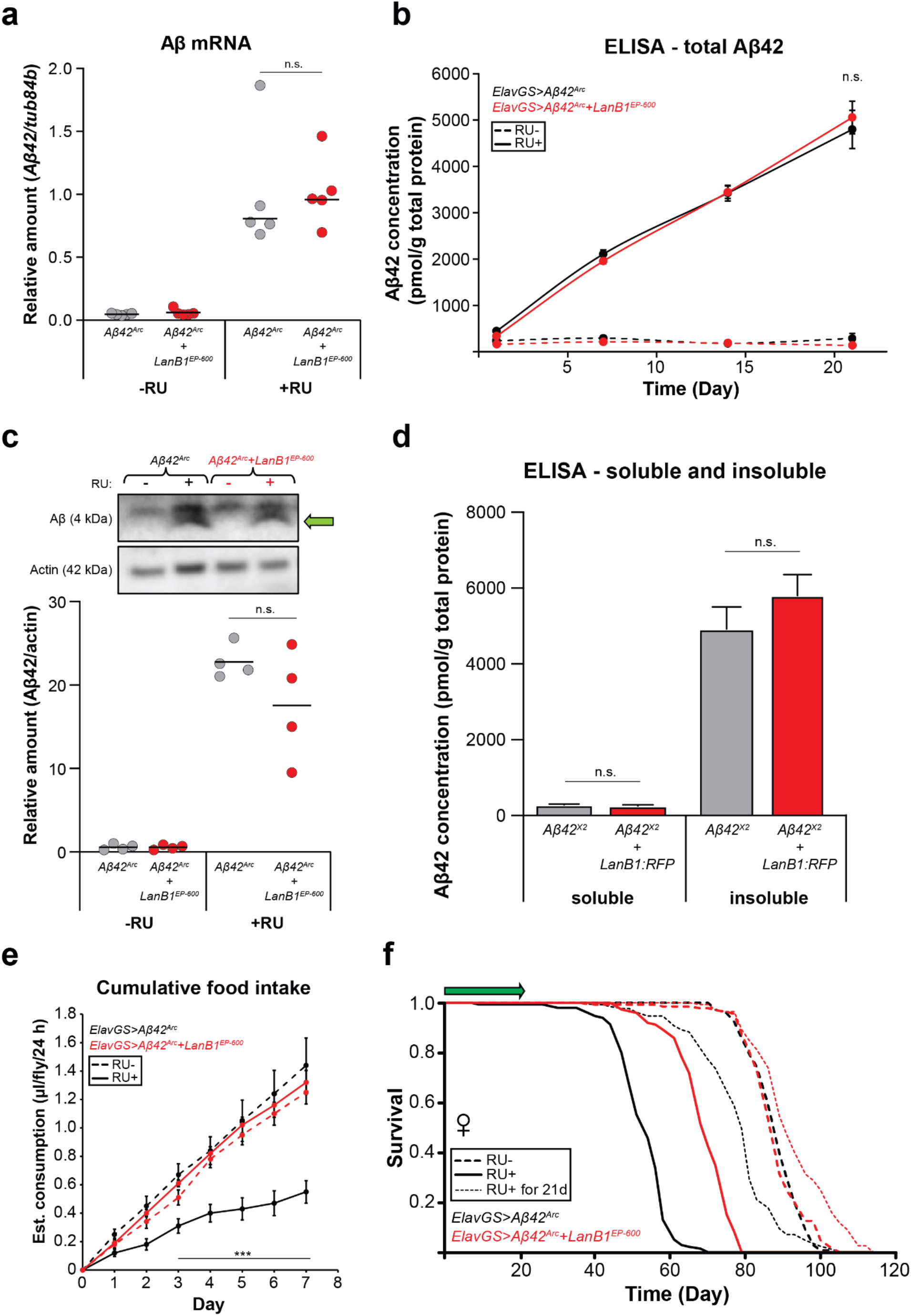
LanB1 did not affect Aβ RNA or protein levels. **a** *Aβ* mRNA was significantly up-regulated upon RU induction (p < 0.0001; one-way ANOVA with Tukey’s post-hoc test) and was not affected by co-expression of LanB1. Data indicate the mean (n = 5 biological replicates per condition). **b** Total Aβ_42_ protein levels increased significantly over time with RU induction compared to uninduced controls (p < 0.0001, two-way ANOVA with Tukey’s post-hoc test), and was unaffected by LanB1 co-expression. Data are shown as mean ± SEM (n = 4 biological replicates per time point). **c** Western blot with quantification of soluble Aβ. Green arrow indicates the distinctive crescent shape of Aβ used for quantification. Soluble Aβ levels increased in induced conditions and was unaffected by LanB1 co-expression. Data indicate the mean (n = 4 biological replicates per condition). **d** After 3 weeks of induction, neither soluble nor insoluble Aβ levels were affected by LanB1 co-expression. Data are shown as mean ± SEM (n = 7 biological replicates per condition). **e** After 3-weeks of RU, induced Aβ^Arc^ flies consumed a significantly smaller amount of food over 7 days compared to uninduced controls (p < 0.0001, two-way ANOVA with Tukey’s post-hoc test). The amount of food ingested by flies with a history of LanB1 and Aβ^Arc^ induction was not significantly different to uninduced controls. Food consumption was measured for 10 flies per condition every 24-hr. Data are shown as mean ± SEM. **e** Expression of Aβ for the first 3 weeks of adulthood significantly reduced lifespan compared to uninduced controls (p = 4.07 x 10^−12^; log rank test). Three-week induction of Aβ and LanB1 together resulted in a small but significant lifespan extension compared to uninduced controls (p = 5.04 x 10^−06^; log rank test).

Expression of Aβ can also have toxic effects during development. For instance, expression of Aβ^X2^ using the GMR-GAL4 driver causes degeneration of the developing eye, resulting in a smaller, glassier appearance compared to the large eye and highly ordered ommatidial lattice in control flies. LanB1 co-expression led to a rescue of both the size and organisation of the eye (**Fig. 1g**). LanB1 could therefore rescue multiple toxic effects of Aβ expression.

### Aβ is not toxic in glia in adult flies

We also induced adult-onset expression of Aβ in the other major cell-type in the fly brain, glia, and examined whether LanB1 co-expression could rescue any deleterious phenotypes. Pan-glial expression of Aβ^Arc^ did not reduce lifespan compared to uninduced controls, nor did Aβ and LanB1 co-expression (**Supplementary Fig. 1e**). This result is in line with the recent finding that Aβ produced by glia was less toxic despite having a much higher brain load than with neuronal expression^23,61^.

### *LanB1*, and not *cdc14*, an adjacent gene in the opposite genomic orientation, is responsible for the rescue of Aβ toxicity

The LanB1^EP-600^ P-element (P{GSV1}) contains UAS sequences at both ends oriented outwards (**Supplementary Fig. 3a**)^62^, and could potentially drive expression of *cdc14,* an adjacent gene in the opposite genomic orientation, previously reported to be involved in stress-resistance and lipid metabolism^63^. To determine if the expression of *cdc14* was affected we performed qPCR on the heads of ElavGS>Aβ^Arc^ flies with and without RU-induction and examined expression of *LanB1* and *cdc14*. We confirmed ∼10-fold increase in *LanB1* gene expression when induced in both UAS-LanB1^EP-600^ and UAS-LanB1:RFP flies (**Supplementary Fig. 3b**), and no change in UAS-cdc14^EY10303^ flies. As predicted, *cdc14* expression was significantly up-regulated in both UAS-cdc14^EY10303^ and UAS-LanB1^EP-600^ flies, but not in UAS-LanB1:RFP transgenic flies (**Supplementary Fig. 3c**).

To determine if LanB1, cdc14 or both were protective against Aβ toxicity, we expressed them singly and examined the effect on eye phenotypes. Up-regulation of cdc14 alone during development, using UAS-cdc14^EY10303^, did not rescue the toxic effects of Aβ^X2^, while UAS-LanB1:RFP rescued both the size and organisation of the eye to the same extent as in UAS-LanB1^EP-600^ flies (**Supplementary Fig. 3d** and quantified in **3e**). We also tested whether cdc14 alone could rescue the short lifespan of ElavGS>Aβ^Arc^ flies. Up-regulation of cdc14 caused a small rescue of Aβ toxicity compared to induced ElavGS>Aβ^Arc^ controls, although a similar small rescue was observed with cytoplasmic GFP-overexpression (**Supplementary Fig. 3f**). LanB1 up-regulation, via either UAS-LanB1^EP-600^ or UAS-LanB1:RFP, caused equivalently pronounced rescue of Aβ toxicity. Thus, LanB1 and not cdc14 was responsible for the rescue of Aβ toxicity in UAS-LanB1^EP-600^ flies.

### Co-expression of LanB1 does not lower levels of Aβ

A potential explanation for the rescue of Aβ toxicity by LanB1 is that less Aβ transcript was generated due to GAL4 titration by the second UAS-transgene. We therefore measured Aβ mRNA and protein levels. Aβ was increased >5-fold when induced in either the presence or absence of the LanB1 transgene (**Fig. 2a**), ruling out GAL4 titration as a factor in the rescue of Aβ toxicity. We then examined the dynamics of Aβ peptide accumulation over a 3-week period by ELISA and found no difference between flies expressing Aβ alone and those co-expressing LanB1 (**Fig. 2b**). We then measured the levels of soluble and insoluble Aβ by western blot (**Fig. 2c**) and ELISA (**Fig. 2d**). There was no significant difference in the levels of soluble or insoluble Aβ between flies expressing Aβ alone and those co-expressing LanB1. Thus, there was no difference in Aβ expression at either the RNA or protein level between flies expressing Aβ alone and those co-expressing LanB1, and nor was the solubility state of Aβ affected, indicating that LanB1 rescued Aβ toxicity rather than Aβ expression.

### Rescue of Aβ toxicity by acute LanB1 over-expression implicates soluble Aβ as the main driver of toxicity

Experiments where expression of Aβ is temporarily induced have shown that high levels of insoluble Aβ persist in the brain long after induction has ceased, while soluble Aβ appears to be rapidly cleared^64^. Induction of Aβ in the first weeks after eclosion results in reduced climbing ability and survival^64^. It remains unclear if the detrimental effects of Aβ on climbing and lifespan in later-life are caused by long-lasting toxic effects of soluble Aβ during the early-life induction period, or from the insoluble Aβ that persists in the ageing brain, or both. We found that 21 days of Aβ induction resulted in a significantly reduced lifespan compared to uninduced controls (**Fig. 2f**), although the reduction was not as great as with chronic Aβ induction. Co-expression of both Aβ and LanB1 for 21 days, in contrast, resulted in a lifespan extension compared to uninduced controls (**Fig. 2f**). These results indicate that the rescue occurred during the early induction period, and that therefore soluble Aβ during the induction phase, rather than effects of accumulated insoluble Aβ, may have been responsible for the later effect on lifespan.

### Laminin accumulates in the ER

The laminin heterotrimer is a canonical ECM protein and is secreted into the extracellular space^65^. Over-expression of monomeric RFP-tagged LanB1, however, leads to intracellular retention^25^. To determine the cellular localization of LanB1, adult brains expressing Aβ and LanB1:RFP were dissected, stained and imaged. In agreement with previous studies^18,55,61^, we found that over-expression of the Aβ^Arc^ peptide resulted in aggregation of Aβ in neurons (**Fig. 3a**). Aβ did not overlap with the nuclear protein histone H3. LanB1 accumulated intracellularly, and partially overlapped with Aβ (**Fig. 3b**).

**Fig. 3.**
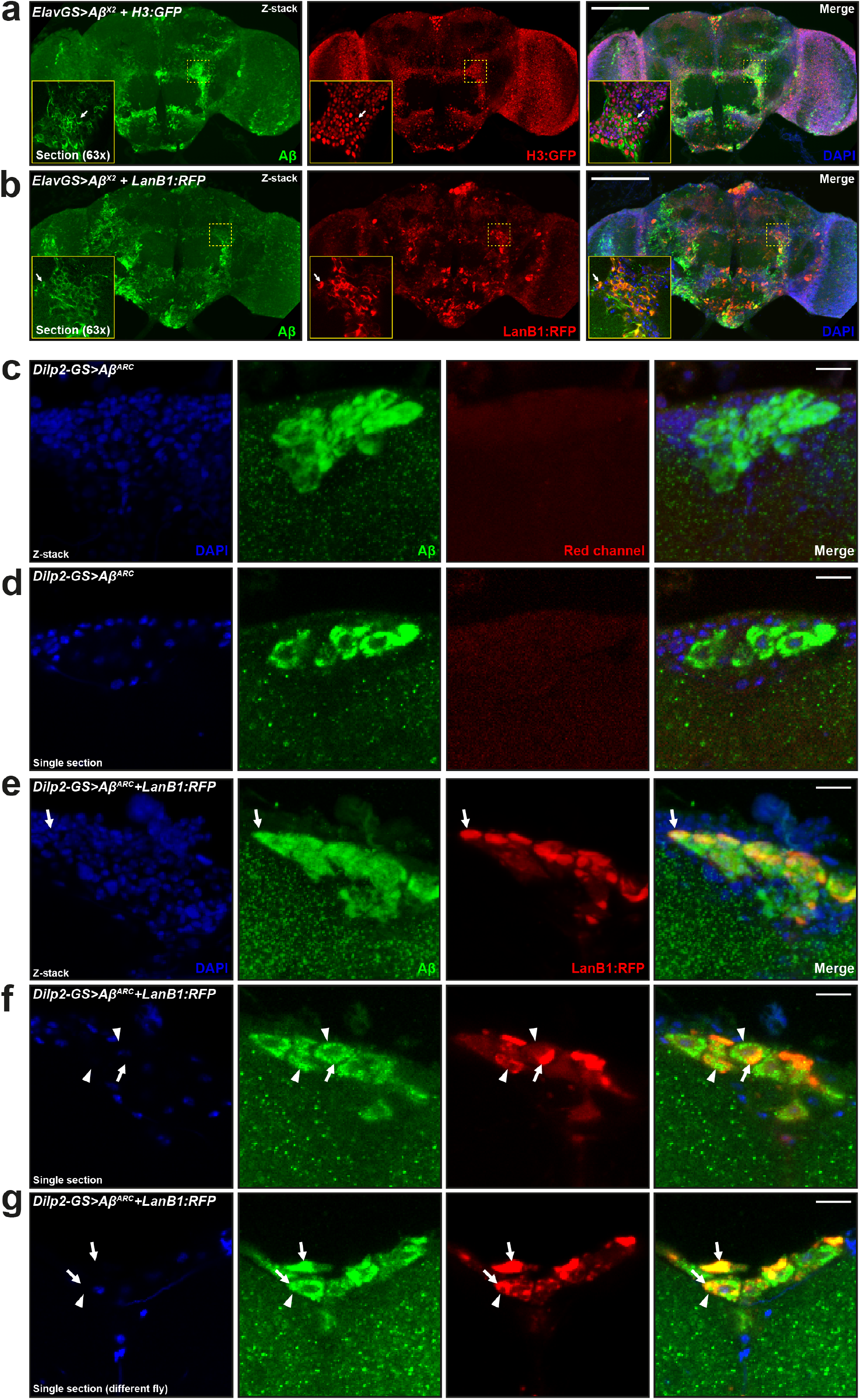
Intracellular accumulation of Aβ and LanB1. **a** Aβ accumulated intracellularly and did not co-localise with the nuclear marker histone H3:GFP. Arrow indicates non-nuclear expression of Aβ. **b** Similarly, LanB1 accumulated intracellularly, not in the nucleus, and partially overlapped with Aβ expression. Arrow indicates co-localisation of Aβ and LanB1. **a,b** Representative confocal fluorescence z projections taken at 20x magnification of whole brains from 21-day-old female flies stained with Aβ (6E10 – green) and DAPI (blue). Yellow box inset shows a single section of the same brain taken at 63x magnification. Endogenous fluorescence (i.e. without staining) of H3:GFP and LanB1:RFP is shown. H3:GFP has been false-coloured red to aid comparison. **c** Z-projection and **d** single section showing intracellular accumulation of Aβ in Dilp2 neurons. **e** Z-projection and **f,g** single sections showing intracellular accumulation of Aβ and LanB1 in Dilp2 neurons. LanB1 expression was not uniformly distributed through the cytoplasm of these cell bodies and appeared to accumulate in discrete intracellular compartments. Arrows highlight co-localisation of Aβ and LanB1, while arrowheads highlight areas with no overlap, often in the same cell. **c-g** Representative confocal images taken at 63x magnification of Dilp2 neurons from 21-day-old female flies stained with Aβ (6E10 – green) and DAPI (blue). Endogenous fluorescence (i.e. without staining) of LanB1:RFP is shown. Genotypes: **a**, *ElavGS>Aβ^X2^ + H3:GFP*; **b**, *ElavGS>Aβ^X2^ + LanB1:RFP*; **c,d**, *Dilp2-GS>Aβ^Arc^*; **e-g**, *Dilp2- GS>Aβ^Arc^ + LanB1:RFP*. Scale bar, 100 μm for **a,b**, 10 μm for **c-g**.

Upon closer inspection, LanB1 appeared to accumulate in discrete intracellular puncta/compartments. To verify the intracellular expression of LanB1 in neurons, we used the inducible Dilp2-GS driver, which drives expression in the Dilp2 neurons. *Drosophila* insulin-like peptide 2 (Dilp2) neurons consist of 10-14 insulin-like peptide producing cells located in the dorsal brain *pars intercerebralis* that project their axons ventrally^66^. Driving LanB1 expression in these neurons allowed us to observe whether or not LanB1 was secreted into the extracellular space and/or accumulated intracellularly. We verified that Aβ and LanB1 were induced in these neurons only when the flies were fed RU486 (**Supplementary Fig. 4**) and that Aβ in Dilp2 neurons accumulated predominantly in the soma (**Fig. 3c,d**). Similar to Aβ, LanB1 accumulated intraneuronally but did not completely overlap with Aβ (**Fig. 3e-g**). LanB1 was not uniformly distributed through the cytoplasm of these cell bodies and appeared to accumulate in discrete intracellular compartments.

As laminin β and γ subunits cannot be secreted from the ER without first forming a heterotrimer^67,68^, over-expressing LanB1 could have led to its accumulation in the ER^53^. Indeed, intracellular accumulation of LanB1 protein has been previously reported for expression of the LanB1:RFP transgene in fat body^25^. Similarly, the expression of the LanB1^EP-600^ element also led to intracellular laminin accumulation^53,69^. To test if the intracellular accumulation of LanB1 was in the ER, we co-expressed LanB1:RFP and a GFP-tagged ER marker, KDEL:GFP, in adult neurons. We found a strong overlap of KDEL:GFP and LanB1:RFP expression, indicating that LanB1 accumulated in the ER (**Fig. 4**). We also found intracellular accumulation, probably in the ER, of LanB2:GFP in Dilp2 neurons when we examined these flies by live 2-photon imaging (**Supplementary Fig. 6**).

**Fig. 4.**
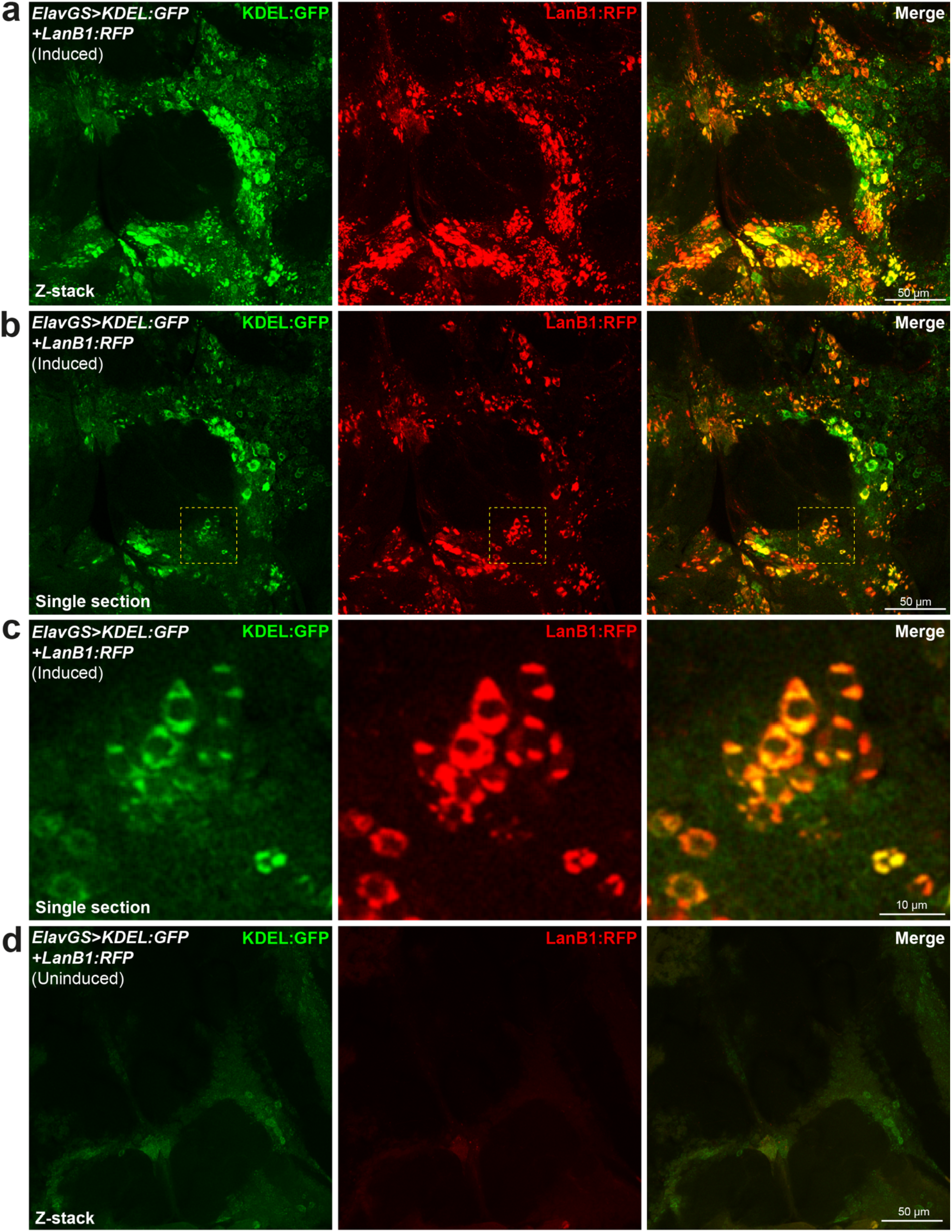
LanB1 accumulates in the ER. **a** Z-projection and **b** single section showing neuronal expression of LanB1 with the ER marker, KDEL:GFP, using the ElavGS driver. **c** Magnified view of the dashed yellow box inset in **b**. LanB1 colocalizes with KDEL:GFP. **d** without RU486 induction, there was very little induction of LanB1 and/or KDEL:GFP expression. Representative confocal fluorescence images taken at 63x magnification from 7-day-old female flies. Images show the antennal lobe and surrounding brain regions. Endogenous fluorescence (i.e. without staining) of KDEL:GFP and LanB1:RFP is shown. Genotype: *ElavGS>KDEL:GFP + LanB1:RFP*. Scale bar, 50 μm for **a,b,d**, 10 μm for **c**.

To determine whether intracellular Aβ aggregates were localized to the ER, we co-expressed Aβ and KDEL:GFP in adult Dilp2 neurons. Aβ did not colocalize with KDEL:GFP, indicating that Aβ did not accumulate in the ER (**Supplementary Fig. 7**). These results are in line with previous studies in adult fly brains that demonstrated that Aβ mainly accumulates in lysosomes^70,71^, and it also does so in mice expressing intraneuronal Aβ^72^.

### LanB1 does not reduce secretion of Aβ from neurons

Since Aβ protein levels, both soluble and insoluble, were not reduced in LanB1 co-expressing flies, we hypothesised that the intracellular accumulation of LanB1 might have prevented the normal secretion of the toxic Aβ peptide into the extracellular space, despite the presence of a signal peptide. We used the Dilp2-GS line to drive expression during the first 3 weeks of adulthood in Dilp2 neurons, and then measured Aβ fluorescence in the area outside of Dilp2-expressing cells. We found that, when induced, Aβ was highly expressed in the cell bodies of Dilp2 neurons and also displayed a diffuse punctate pattern of staining, compared to very little staining in uninduced flies (**Fig. 5a**). When quantified, we found a significant increase in Aβ fluorescence in induced flies compared to uninduced controls, indicating that Aβ is secreted into the extracellular environment (**Fig. 5b**). LanB1 co-expression had no effect on secretion of Aβ. In summary, Aβ aggregated strongly in the cell bodies of Dilp2 neurons and was also secreted into the extracellular milieu (**Fig. 5c**). Therefore, the beneficial effects of LanB1 co-expression were not due to the prevention of Aβ secretion.

**Fig. 5.**
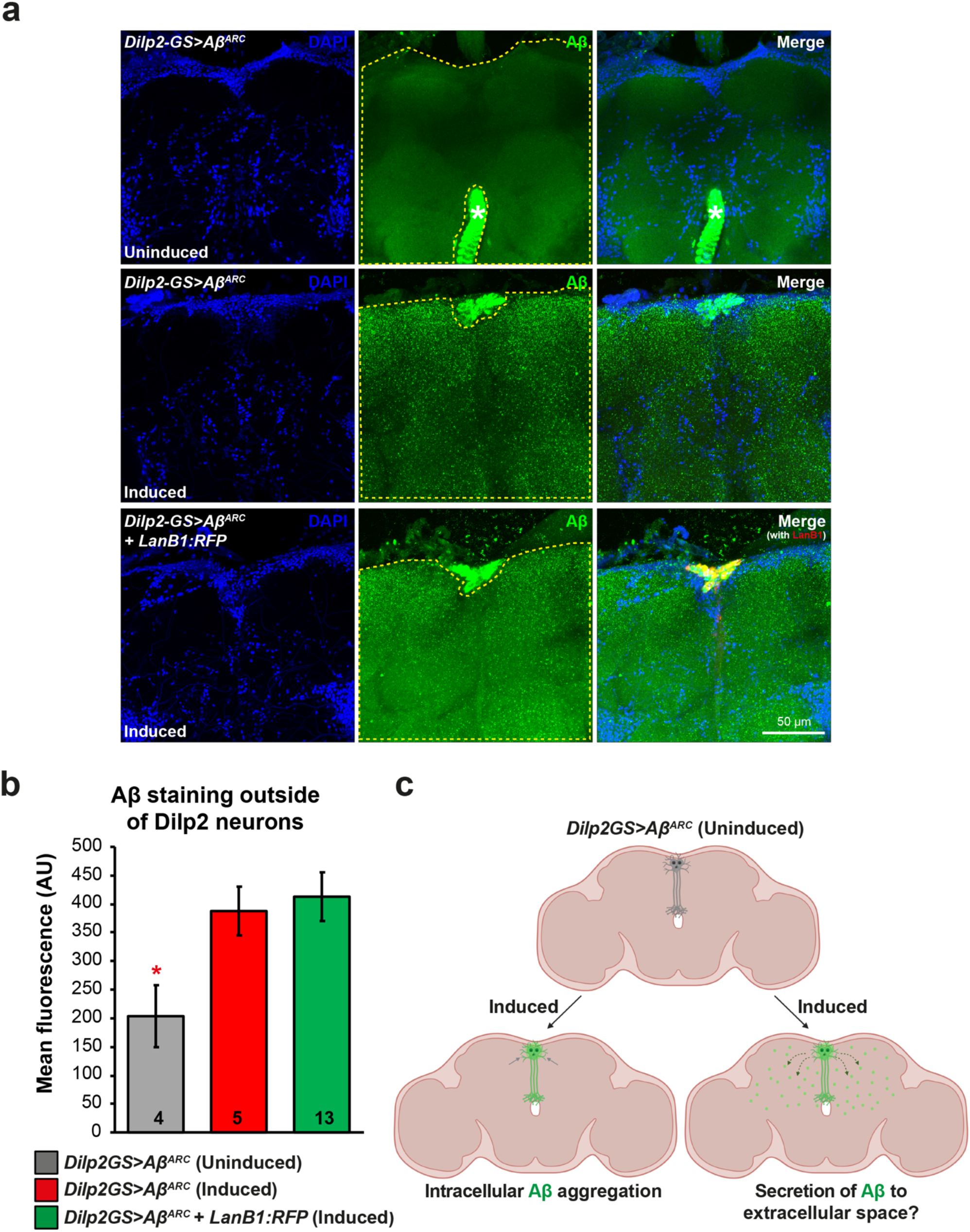
Quantification of Aβ secretion from Dilp2 neurons. **a** Aβ expression in Dilp2 neurons was highest in the cell body area, but there was also diffuse, punctate staining of Aβ outside the cell bodies. LanB1 co-expression had no effect on Aβ fluorescence outside Dilp2 neurons. Without RU induction, there was no Aβ found in Dilp2 neurons or the surrounding area. Representative confocal fluorescence z projections taken at 63x magnification of whole brains from 21-day-old female flies stained with Aβ (6E10 – green) and DAPI (blue). Dashed yellow area indicates the area of fluorescence measurement. White asterisk in the top row of images indicates strong staining of oesophageal muscle. Endogenous fluorescence (i.e. without staining) of LanB1:RFP is also shown. **b** Quantification of Aβ fluorescence outside Dilp2 neurons. There was a significant increase in diffuse, punctate Aβ staining outside Dilp2 neurons when induced (p = 0.048; one-way ANOVA). LanB1 co-expression had no effect on Aβ staining. Biological replicate numbers for each condition are labelled within the bars. **c** Diagram of the proposed secretion of Aβ from Dilp2 neurons into the extracellular space. Created with BioRender.com. Genotypes: *Dilp2-GS>Aβ^Arc^*; and *Dilp2-GS>Aβ^Arc^ + LanB1:RFP*. Scale bar, 50 μm.

### Increased expression of a Collagen IV subunit also rescues Aβ toxicity in adult neurons

The trimerisation of laminins takes place in the ER and all three subunits are required for secretion to occur^49^. We hypothesized that over-expressing protein subunits of similar, large, obligate heterotrimers that assemble in the ER may lead to amelioration of Aβ toxicity. Collagen IV, the main component of basement membranes, is another obligate heterotrimeric protein formed by three α chains (two α chains of Collagen at 25C (Cg25C) and one α chain Viking (Vkg)), and is present in all metazoans^73,74^. Similar to laminins, expression of collagen IV subunits in the adult fly brain was predominantly in haemocytes and a subset of glial cells, while endogenous expression in neurons was low (**Supplementary Fig. 2c**), indicating that pan-neuronal expression of these subunits by ElavGS or NsybGS is ectopic. The RFP-tagged Cg25C transgenic over-expression line (*UAS-Cg25C:RFP*) rescued the short lifespan from induction of Aβ^Arc^ (**Fig. 6a**) and Aβ^X2^ (**Fig. 6b**), and to the same degree as LanB1, potentially indicating a similar mechanism of action.

**Fig. 6.**
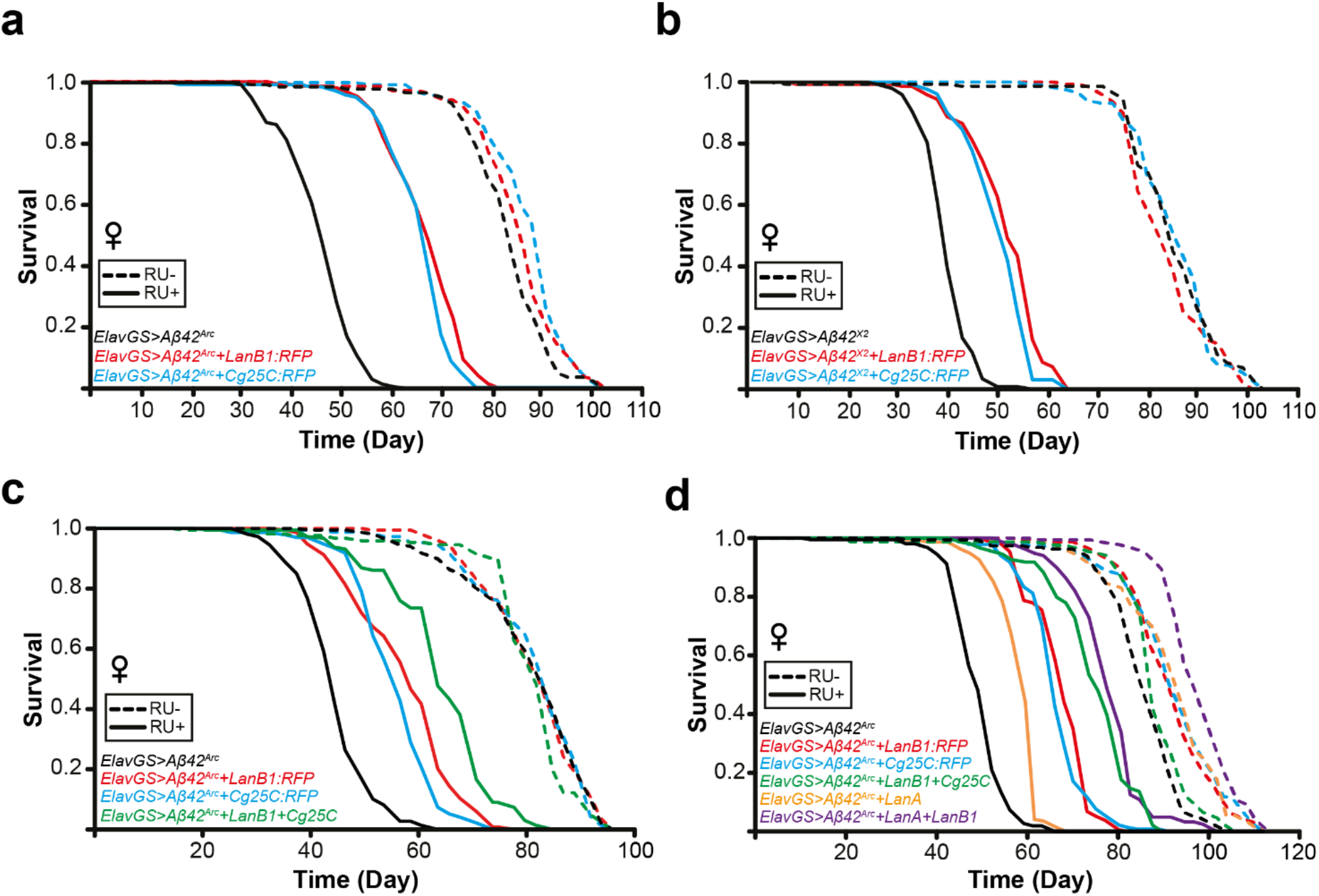
Enhanced rescue of Aβ toxicity with combined laminin/collagen IV subunit over-expression. **a** LanB1 significantly rescued Aβ toxicity (p = 2.85 x 10^−62^; log rank). Cg25C significantly rescued Aβ toxicity (p = 3.42 x 10^−63^; log rank). There were small but significant extensions of lifespan in the uninduced controls (LanB1, p = 0.036; Cg25C, p = 7.51 x 10^−06^; log rank vs *ElavGS>Aβ^Arc^* alone). **b** LanB1 significantly rescued Aβ toxicity (p = 8.04 x 10^−40^; log rank). Cg25C significantly rescued Aβ toxicity (p = 1.25 x 10^−35^; log rank). **c** LanB1 significantly rescued Aβ toxicity (p = 3.77 x 10^−33^; log rank). Cg25C significantly rescued Aβ toxicity (p = 2.24 x 10^−30^; log rank). LanB1 + Cg25C had a partially additive effect on the rescue of Aβ^Arc^ toxicity. Cox Proportional Hazard analysis showed a significant interaction between LanB1 and Cg25C (p < 0.001). **d** LanB1, Cg25C, and LanA rescued Aβ toxicity (LanB1, p = 3.34 x 10^−65^; Cg25C, p = 2.11 x 10^−55^; LanA, p = 1.10 x 10^− 32^; log rank vs *ElavGS>Aβ^Arc^* alone). LanB1 + Cg25C, and LanB1 + LanA had a partially additive effect on the rescue of Aβ^Arc^ toxicity. Cox Proportional Hazard analysis showed a significant interaction with LanB1 for Cg25C (p < 0.001) and LanA (p = 0.022). There was significant extension of lifespan in the uninduced controls (LanB1, p = 6.45 x 10^−10^; Cg25C, p = 1.57 x 10^−11^; LanA, p = 1.40 x 10^−10^; LanA+LanB1, p = 1.66 x 10^−39^; LanB1+Cg25C, p = 0.026; log rank vs *ElavGS>Aβ^Arc^* alone). Dashed lines represent uninduced ‘RU-‘ controls, solid lines represent induced ‘RU+’ conditions. For all lifespan experiments n = 150 flies per condition.

We next examined if the rescue of degeneration in the fly eye due to Aβ toxicity (**Fig. 1f**) was specific to LanB1. Co-over-expression of mCD8:GFP or mCD8:RFP alone, as controls for GFP/RFP transgene overexpression, had no significant effect on Aβ toxicity (**Supplementary Fig. 5a** and quantified in **5b**). First, we assessed the effect of modulation of other laminin/collagen subunits on the rough eye phenotype. We confirmed that LanB1 co-expression (either using *UAS-LanB1^EP-600^* or *UAS-LanB1:RFP*) significantly rescued Aβ toxicity. Additionally, we found that co-expression of laminin subunits, LanA (*UAS-LanA^EY02207^*) and LanB2 (*UAS-LanB2:GFP*), and the collagen IV subunit Cg25C (*UAS-Cg25C:RFP*) separately all rescued Aβ toxicity (**Supplementary Fig. 5**). Knockdown of laminin α-chain subunits, LanA and wb, using TRiP RNAi transgenics had no effect on toxicity, while knockdown of γ-chain LanB2 led to a small but significant rescue (**Supplementary Fig. 5**). Next we generated recombinant transgenic lines for co-expression of LanB1, i.e. ‘LanB1^EP-600^ + Cg25C:RFP’, ‘LanB1^EP-600^ + LanA^OE^’, ‘LanB1^EP-600^ + LanB2^TRiP^’, and ‘LanB1^EP-600^ + wb^TRiP^’. Although Cg25C and LanA could individually rescue eye size, modulation of these other subunits had no effect on the ability of LanB1 to protect against Aβ toxicity, indicating that LanB1 was the primary cause of the toxicity rescue in the context of the developing eye (**Supplementary Fig. 5**).

### Enhanced rescue of Aβ toxicity with combination of laminin and collagen subunits

As both LanB1 and Cg25C individually rescued Aβ toxicity to a similar extent (**Fig. 6a,b**), and the combination of both did not further rescue Aβ toxicity in the developing eye (**Supplementary Fig. 5**), we hypothesized that these proteins may have been epistatic, i.e. acting via the same mechanism. We therefore examined the effect of the combination of LanB1 and Cg25C in adult neurons on lifespan. We found that LanB1 and Cg25C together resulted in a greater rescue of Aβ toxicity compared to individual co-expression (**Fig. 6c**). Cox Proportional Hazard analysis showed a significant interaction between LanB1 and Cg25C, indicating that, although the enhanced rescue was partially additive, it was also acting via a shared pathway.

Laminin A (LanA) is one of two laminin α-chains in *Drosophila*, the other being *wing blister* (*wb*), which form heterotrimers with LanB1 (β-chain) and LanB2 (γ-chain)^48^. We found that over-expression of LanA alone could significantly rescue Aβ toxicity, but not to the same degree as LanB1 or Cg25C alone (**Fig. 6d** and **Supplementary Fig. 8a**). We generated double transgenic flies with both LanA and LanB1, which resulted in an even greater rescue of toxicity, indicating an additive effect. However, Cox Proportional Hazard analysis showed a significant interaction between LanB1 and LanA, indicating the enhanced rescue is partly acting via a shared pathway. LanA also rescued Aβ toxicity in the developing fly eye when compared to Aβ^X2^ alone, though not to the same extent as LanB1 (**Supplementary Fig. 5**). These results indicate that LanA can rescue Aβ toxicity, but to a lesser degree than LanB1. Overall, the collagen IV subunit Cg25C and the laminin α chain LanA led to enhanced rescue of Aβ toxicity in combination with the laminin β chain LanB1. These effects were epistatic, suggesting an overlapping molecular pathway.

### LanB1 rescue of Aβ toxicity is independent of the BiP/Xbp1 ER stress response pathway

Aβ induces ER stress markers, including the ER chaperone BiP, and leads to increased alternative splicing of Xbp1^21–23^. Increased BiP can exacerbate Aβ toxicity while reduction of BiP has been shown to have beneficial effects^22,75^. To assess if BiP reduction was behind the LanB1 rescue of Aβ toxicity, we examined BiP mRNA and protein levels. After 3-weeks of Aβ induction, BiP was significantly up-regulated at both the mRNA (**Fig. 7a**) and protein level (**Fig. 7b,c**) in fly heads. Co-expression of LanB1 with Aβ had no effect on BiP mRNA or protein levels. Therefore, LanB1 rescued Aβ toxicity independently of the ER stress regulator, BiP.

**Fig. 7.**
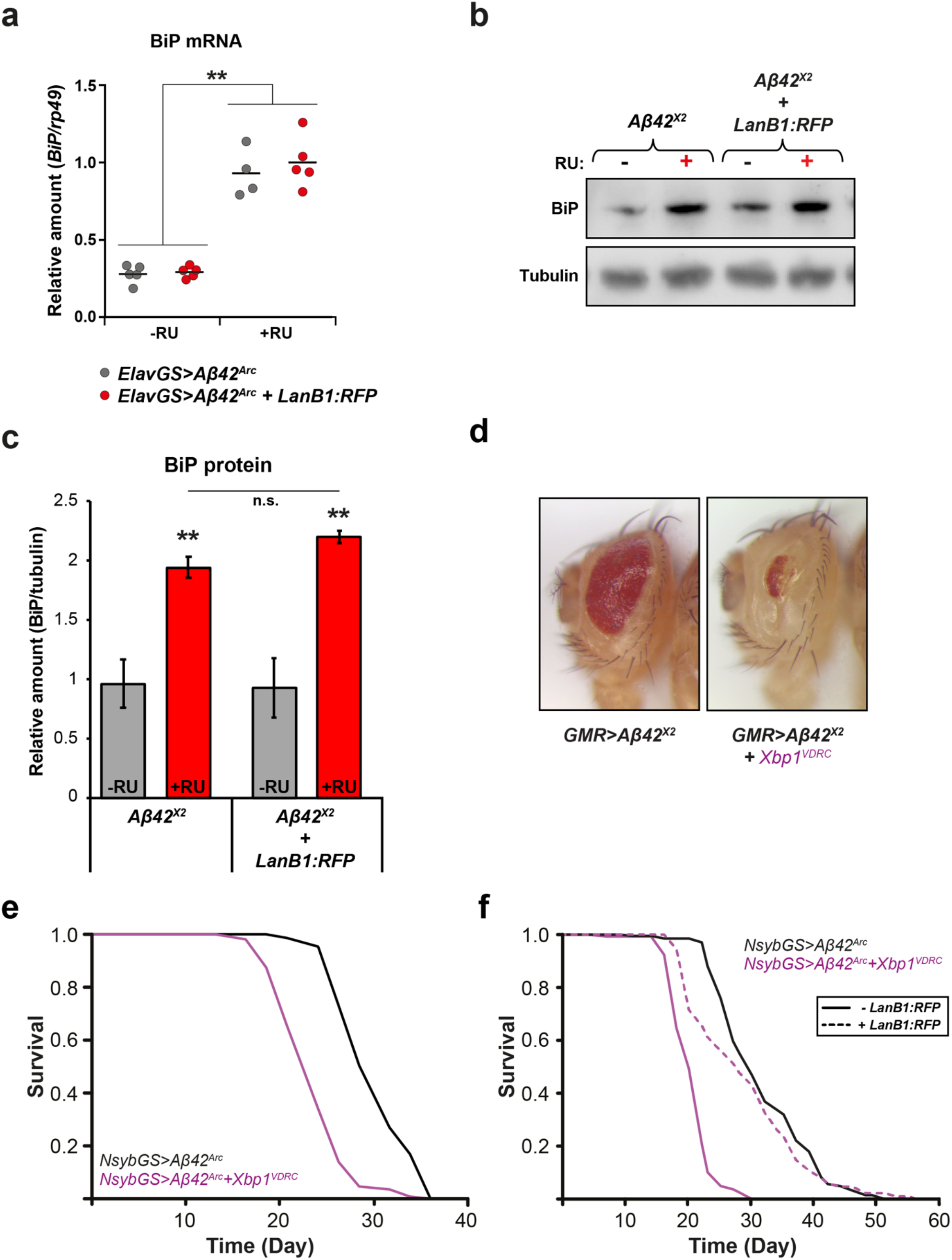
LanB1 rescue of Aβ toxicity is independent of the BiP/Xbp1 ER stress response pathway. **a** *BiP* mRNA was significantly upregulated upon Aβ induction (p < 0.01; one-way ANOVA with Tukey’s post-hoc test) and this was not changed by the overexpression of LanB1:RFP. Data are shown indicating the mean (n = 4-5 biological replicates per condition). **b** Western blot of BiP protein levels, and quantification in **c** confirm that BiP protein is also significantly upregulated with Aβ induction and this is not changed by LanB1:RFP overexpression (p < 0.01; one-way ANOVA with Tukey’s post-hoc test). Data are shown indicating the mean (n = 3-4 biological replicates per condition). **d** Co-expression of Aβ^X2^ with Xbp1^VDRC^ in the developing eye. Knockdown of Xbp1 significantly exacerbated Aβ toxicity and these flies exhibited very small and depigmented eyes. **e** Knockdown of Xbp1 exacerbated Aβ toxicity and significantly shortened lifespan compared to controls (p = 3.03 x 10^−30^; log rank). **f** Knockdown of Xbp1 exacerbated Aβ toxicity and significantly shortened lifespan compared to controls (p = 1.68 x 10^−44^; log rank). Over-expression of LanB1 rescued the shorter lifespan of Xbp1^VDRC^ (Aβ+LanB1:RFP+Xbp1^VDRC^ vs Aβ+Xbp1^VDRC^, p = 9.13 x 10^−22^; log rank). For display purposes, the control micrograph in **d** is the same as that in **Supplementary Fig. 5**. For all lifespan experiments n = 150 flies per condition. ‘VDRC’ indicates an RNAi transgene. Genotypes: **a**, *ElavGS>Aβ^Arc^*; *ElavGS>Aβ^Arc^ + LanB1:RFP*; **b,c**, *NsybGS>Aβ^Arc^*; *NsybGS>Aβ^Arc^ + LanB1:RFP*.

An increase in active spliced Xbp1, a marker of IRE1α activation, can ameliorate Aβ toxicity, while a decrease exacerbates Aβ toxicity^21,23^. To assess whether the LanB1 rescue was acting via Xbp1, we examined the effect of Xbp1-knockdown on Aβ toxicity. We used the Aβ-induced rough eye phenotype, and Aβ-induced short-lifespan as read-outs. Knockdown of Xbp1 significantly exacerbated Aβ toxicity in the developing eye (**Fig. 7d**), and also led to a significantly shortened lifespan compared to Aβ alone (**Fig. 7e,f**). Over-expression of LanB1 significantly rescued the short lifespan of Aβ flies with Xbp1-knockdown (**Fig. 7f**), and there was no significant difference in lifespan between these rescue flies and Aβ-alone controls (p = 0.067, log-rank). Thus, the ability of LanB1 to rescue Aβ toxicity was independent of Xbp1. Since LanB1 did not affect BiP levels in the context of neuronal Aβ, and since LanB1 could rescue Aβ toxicity independently of Xbp1, LanB1 must have acted independently of the IRE1α/XBP1s arm of the ER stress response pathway.

### Proteins that are retained in the ER can rescue Aβ toxicity

Since LanB1 over-expression resulted in intra-ER accumulation, we investigated whether over-expression of other proteins restricted to the ER could result in the same phenotype. To test this, we compared the effects of GFP localised to different cellular compartments, namely membrane-targeted mCD8:GFP, LanB2:GFP (laminin γ chain) which accumulates in the ER, KDEL:GFP which is retained in the ER, and secr:GFP which is secreted extracellularly, and measured their effect on Aβ^X2^ toxicity in the developing eye. As with LanB1 over-expression, expression of LanB2:GFP rescued Aβ toxicity (**Fig. 8a** and quantified in **8b**) compared to mCD8:GFP controls. Expressing KDEL:GFP led to a rescue of eye size while secr:GFP exacerbated toxicity. In adult neurons LanB2 significantly rescued the short lifespan induced by Aβ^Arc^ toxicity (**Fig. 8c**). Again, KDEL:GFP rescued Aβ^Arc^ toxicity and secr:GFP exacerbated Aβ^Arc^ toxicity (**Fig. 8c**). LanB2 also significantly rescued the short lifespan with Aβ^X2^ induction (**Supplementary Fig. 8b**). Thus, the retention of over-expressed proteins in the ER ameliorated Aβ toxicity. Overall, these results suggest that ER-retention is a promising avenue of AD research.

**Fig. 8.**
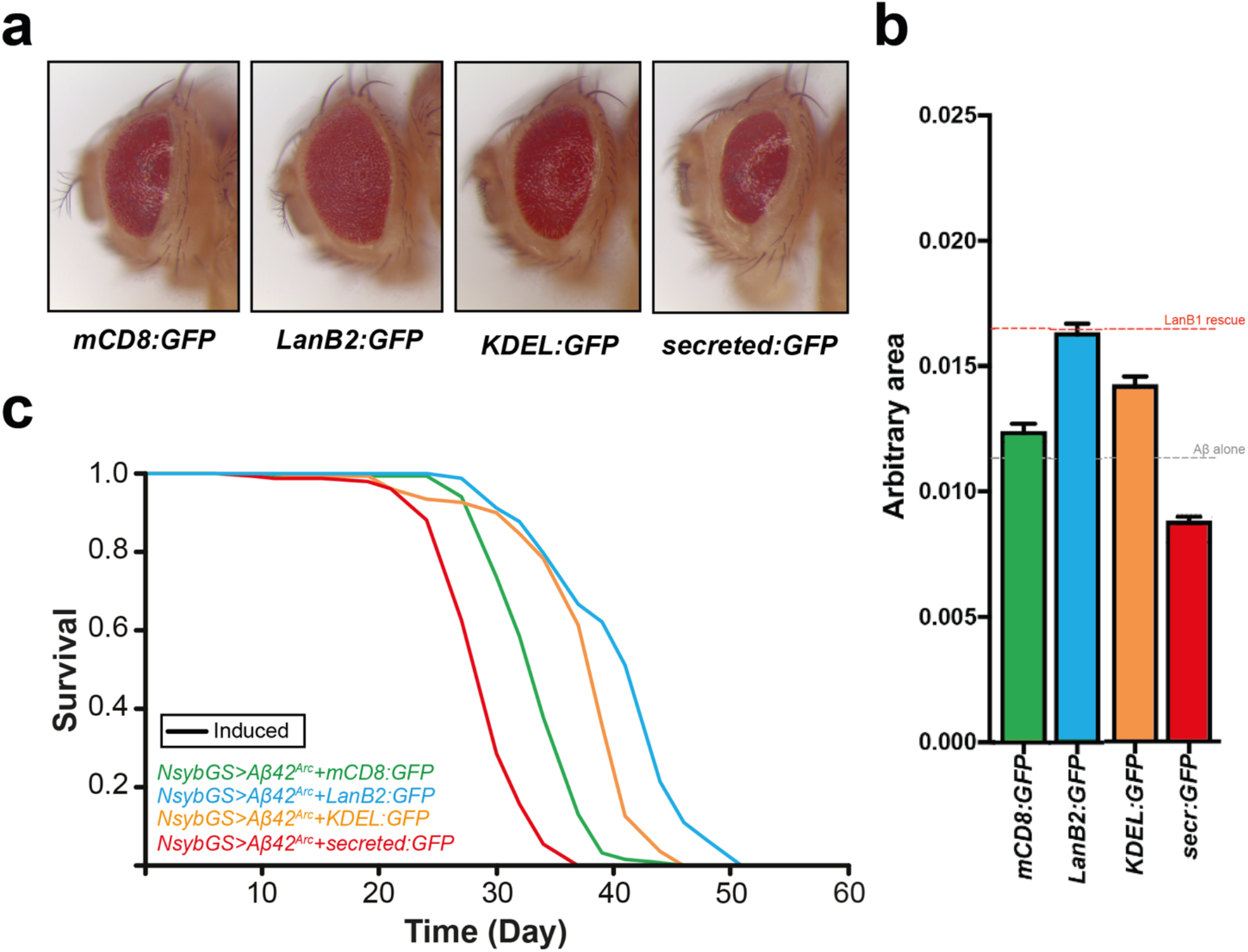
Protein accumulation in the ER may be responsible for the rescue of Aβ toxicity. **a,b** Co-expression of Aβ^X2^ with LanB2:GFP or KDEL:GFP rescued Aβ toxicity in the developing eye compared to mCD8:GFP controls, while secr:GFP exacerbated Aβ toxicity. **b** Quantification of eye sizes in **a**. mCD8:GFP did not rescue the rough eye phenotype compared to Aβ^X2^ alone (see **Supplementary Fig. 5**). LanB2:GFP and KDEL:GFP significantly rescued eye size while secr:GFP significantly reduced eye size compared to mCD8:GFP controls (p < 0.0001; one-way ANOVA with Dunnett’s post-hoc test). Data are shown as mean ± SEM (n = 34-46 eyes measured per condition). For display purposes, two micrographs in **a** are the same as those in **Supplementary Fig. 5**. **c** Survival curves of female flies induced to express Aβ^Arc^ via NsybGS. LanB2:GFP and KDEL:GFP rescued Aβ toxicity while secr:GFP exacerbated Aβ toxicity compared to mCD8:GFP controls (LanB2:GFP, p = 2.37 x 10^−26^; KDEL:GFP, p = 3.49 x 10^−17^; secr:GFP, p = 1.79 x 10^−19^; log rank). Genotypes: **a,b**, *GMR>Aβ^X2^ + mCD8:GFP*; *GMR>Aβ^X2^ + LanB2:GFP*; *GMR>Aβ^X2^ + KDEL:GFP; GMR>Aβ^X2^ + secr:GFP.* For all lifespan experiments n = 150 flies per condition.

### Lamb1 also accumulates in the ER when overexpressed in mouse neural tissue

Disruption of laminin subunit production results in intracellular accumulation of the remaining subunits^53^. To our knowledge, however, there are no published data examining the over-expression of Laminin subunits and their expression pattern in mammalian models. We used *ex-vivo* mouse organotypic hippocampal slice cultures to transduce mouse Lamb1 using engineered lentivirus. Organotypics were generated from P10 wildtype (C57/BL6J) mouse pups and incubated with mouse Lamb1 (mLamb1 + GFP) recombinant lentivirus or control recombinant lentivirus (GFP only). Cultures were transfected with virus at 14 days *in vitro*, and fixed after a further 14 days. Over-expression of mLamb1 resulted in marked intracellular accumulation compared to controls and did not result in a uniform cytoplasmic distribution, indicating compartmentalisation (**Fig. 9a**). Lamb1 colocalised with the ER marker Calnexin in some areas, indicating retention of Lamb1 in the ER, though there were also areas in these cells with no overlap. Therefore, the ER-retention of over-expressed laminin β chain is a conserved process.

**Fig. 9.**
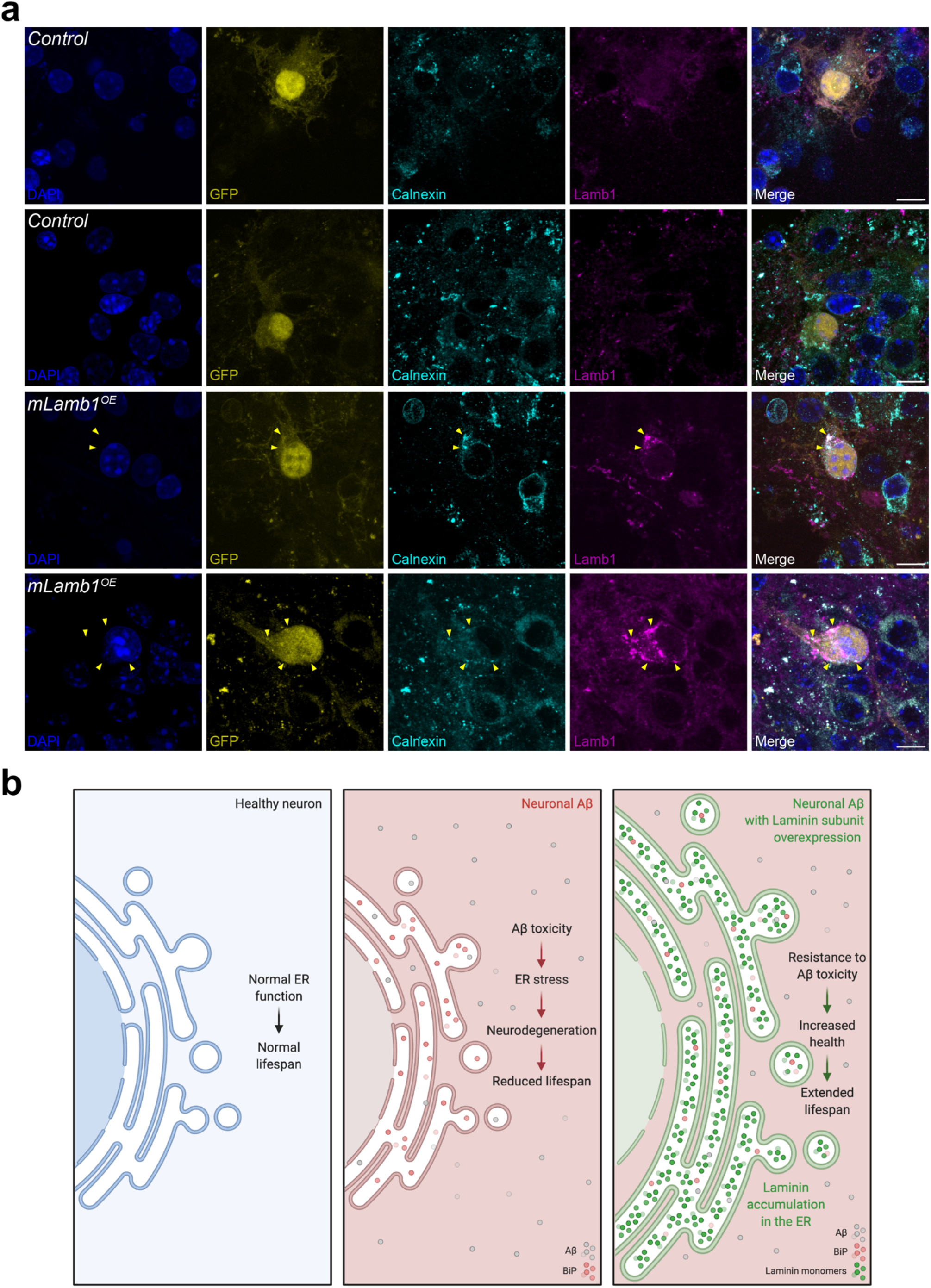
Intra-ER retention of overexpressed Lamb1 is conserved in mouse brain tissue. **a** Organotypic hippocampal slice cultures from 10-day old mouse pups were incubated for 24 hours with lentivirus containing *mLamb1+*GFP or control lentivirus (GFP only), and then fixed 2-weeks later. Slices were then stained for Lamb1 and the ER-marker, Calnexin. Endogenous GFP expression was used to identify successful transduction. DAPI labelled nuclei. Top 2 rows show examples of control lentivirus without *mLamb1* induction. Bottom 2 rows show examples of lentiviral *mLamb1* induction. Yellow arrowheads indicate areas of colocalization of Lamb1 and Calnexin. **b** Summary model of intra-ER laminin retention. Created with BioRender.com. Scale bar, 10 μm.

## Discussion

Here, we showed that ectopic, neuronal over-expression of laminin (and collagen IV) monomers provided robust protection in an *in vivo Drosophila* model of AD, and that ectopic over-expression of LanB1 resulted in ER-retention in these neurons. Over-expression of mouse Lamb1 in *ex vivo* mouse organotypic hippocampal slice cultures also resulted in ER-retention of these monomers, highlighting a conserved process. In the fly, LanB1 rescued Aβ toxicity without reducing Aβ levels (soluble or insoluble) and without affecting Aβ secretion into the extracellular milieu. LanB1 also rescued Aβ toxicity in combination with other laminin subunits and a collagen IV subunit and acted independently of the IRE1α/XBP1s ER stress response branch. Finally, expression of other proteins targeted for retention in the ER could significantly rescue Aβ toxicity. Overall, we have described a novel mechanism whereby ER-retention of proteins, typically detrimental to cellular health, is a potential therapeutic target for AD.

For nearly 3 decades, Aβ has been one of the major targets for AD therapy with most major clinical trials aiming to reduce or detoxify Aβ^76^. Similarly, most studies in *Drosophila* showing rescue of Aβ toxicity appear to work, at least partially, via reduction of Aβ levels^23,56,75,77^. Previous studies in both flies and humans, however, have demonstrated that Aβ load can be uncoupled from toxicity^21,22,78^. Intriguingly, LanB1 was able to substantially rescue Aβ toxicity without altering levels of Aβ, either soluble or insoluble, and also without reducing secretion of Aβ into the extracellular milieu indicating that LanB1 expression increased neuronal resistance to Aβ toxicity. This result also suggests that, since LanB1 accumulated in the ER, the beneficial effect of LanB1 over-expression may also occur in the ER. Further work examining the intracellular/extracellular availability of LanB1 may inform some of the debate surrounding intracellular vs extracellular Aβ toxicity^11^.

Collagen VI rescues Aβ toxicity by sequestering Aβ into large aggregates in the extracellular milieu^35^. Hence, LanB1 could have promoted the formation of Aβ fibrils from the more toxic oligomeric form. However, we observed no difference in the levels of soluble or insoluble Aβ levels with LanB1 co-expression, indicating that LanB1 was not altering the ratio of soluble/insoluble Aβ. It remains to be seen whether Aβ is binding to LanB1, or if LanB1 is sequestering toxic oligomeric Aβ into less toxic structures. Crucially, a rescue of Aβ toxicity was also observed with Cg25C, LanA, LanB2, and KDEL:GFP, the last of which likely does not bind Aβ, indicating mechanisms other than Aβ sequestration by LanB1.

Since glia are the other major cell type in the adult fly brains and can produce laminins, we determined if LanB1 over-expression in these cells could also rescue Aβ toxicity. We could not verify this, however, as we did not observe Aβ toxicity using GliaGS, and co-expression of LanB1 also had no effect. This result was consistent with recent studies that found glial Aβ was less toxic than neuronal Aβ despite having a much higher brain load than those produced by neurons^23,61^.

ER stress has been implicated in the progression of AD^4^, although the precise mechanisms that underlie protein misfolding contributions to AD pathogenesis are still unclear^15^. Here, we found that BiP was not affected at the mRNA/protein level by LanB1 despite the rescue of Aβ toxicity. It is possible that BiP expression was at the upper limit and LanB1 could not increase BiP expression further. Regardless, it is likely that BiP was chronically high throughout adult lifespan, which is detrimental to neuronal health when co-expressed with Aβ^75^. Intriguingly, BiP may be a molecular chaperone of laminin assembly in the ER^79^, which may indicate that LanB1 over-expression sequesters BiP from having toxic effects when upregulated chronically.

The importance of Xbp1 in the response to Aβ is evident with knockdown of Xbp1, which has been shown to enhance Aβ toxicity^21,23^. However, Lanb1 could rescue the enhanced toxicity of Aβ+Xbp1^RNAi^, indicating that Lanb1 could act independently of Xbp1. In addition, knocking down Xbp1 increased Aβ protein levels while over-expression reduced Aβ protein levels^23^. However we found no difference in Aβ levels with LanB1 co-expression, further indicating that the beneficial effects are independent of Xbp1. Further work analyzing epistatic interactions between LanB1 over-expression and the ER stress response branches will be necessary to deduce the mechanism underlying the rescue of Aβ toxicity.

LanA also rescued Aβ toxicity, but not to the same extent as LanB1. Previous studies have shown that the laminin α-chain can be secreted as a monomer without the requirement for heterotrimerisation^49,50^. Thus, it is possible that LanA is secreted from the ER in monomeric form and so the ER may not accumulate as many large proteins, and hence has a reduced ability to rescue Aβ toxicity. If this were the case, it would be expected that over-expression of the other laminin α-chain (wb) might exhibit the same degree of Aβ rescue as LanA. Collagen IV also forms an obligate heterotrimer with 2 chains of Cg25C and one Vkg chain. Similar to LanA, Vkg can be secreted as a monomer^58^ and it is thought that Cg25C requires Vkg for secretion (like LanB1/LanB2). We suspect that over-expression of LanB1/LanB2/Cg25C led to ER retention of these subunits and resulted in ER expansion (illustrated in **Fig. 9b**). We predict that if Vkg can be secreted as a monomer, it may also show a lesser rescue than Cg25C.

Laminin or collagen upregulation is considered detrimental in most contexts^40,80^ and especially in relation to cancer invasiveness^81,82^. Indeed research examining disease-causing mutations in laminin/collagen has focused on defective ECM-receptor signalling as the underlying cause of pathology^80,82^. Other studies have focused on receptor-independent effects of laminin/collagen loss and have found that mutations triggered ER stress, potentially due to defective secretion of remaining subunits^53,67,83^. A case study of a patient with porencephaly (cystic brain lesions) carrying a collagen IV α2 mutation, which caused intracellular accumulation of the COL4A2 chain, found ER stress and UPR activation which could be rescued by chemical chaperone treatment^83^. Interestingly, the patient’s father carrying the same allele also displayed basement membrane defects but no disease symptoms, indicating that intracellular accumulation of COL4A2 rather than extracellular effects led to disease^83^.

Our hypothesis that LanB1 retention in the ER in neurons is beneficial is augmented by the finding that ER-retained GFP (KDEL:GFP) also reduced Aβ toxicity. Interestingly, GFP targeted for extracellular secretion (secr:GFP) exacerbated Aβ toxicity. It is possible that extracellular GFP is degraded by glia and thus reduces the capacity of glia to clear Aβ, leading to increased toxicity. In agreement with our finding that secr:GFP exacerbated Aβ toxicity, Fernandez-Funez *et al*. showed that eyes of GMR>Aβ+secr:GFP flies were smaller when compared to GMR>Aβ+LacZ^84^. In addition, prion protein, a molecule located extracellularly, exacerbated Aβ toxicity when co-expressed, and caused massive Aβ deposition in the brain^85^.

We have discovered that increased expression of ER-retained proteins, typically seen as detrimental, rescued Aβ toxicity in neurons, and could be a new therapeutic avenue for AD research.

## Methods

### *Drosophila* stocks and fly husbandry

The wild-type stock Dahomey was collected in 1970 in Dahomey (now Benin) and has since been maintained in large population cages with overlapping generations on a 12L:12D cycle at 25°C. The white Dahomey (w^Dah^) stock was derived by incorporation of the w^1118^ deletion into the outbred Dahomey background by successive backcrossing.

The following stocks were obtained from the Bloomington *Drosophila* Stock Center: UAS-LanB1^EP-600^ (#43428), UAS-LanA^EY02207^ (#20141), UAS-cdc14^EY10303^ (#16450), UAS-Col13A1^EY09983^ (#17628), GMR-GAL4 (#9146), UAS-GFP (#1521), UAS-H3:GFP (#68241), UAS-mCD8:GFP (#5137), UAS-mCD8:RFP (#27392), UAS-KDEL:GFP (#9898). The following stocks were obtained from the Vienna *Drosophila* Resource Center: UAS-Mys^VDRC^ (β1-integrin) (#29619), UAS-Mew^VDRC^ (αPS1-integrin) (#44890), UAS-wb^VDRC^ (wb^RNAi 1^) (#3141), UAS-Xbp1^VDRC^ (#109312). The following stocks were obtained from the Transgenic RNAi Project at Harvard Medical School: UAS-LanA^TRiP^ (#28071), UAS-LanB2^TRiP^ (LanB2^RNAi 1^) (#55388), UAS-LanB2^TRiP^ (LanB2^RNAi 2^) (#62002), UAS-wb^TRiP^ (wb^RNAi 2^) (#29559). UAS-Aβ_42_ was a gift from Dr. Damian Crowther (University of Cambridge, UK)^55^. UAS-Aβ_42_ (tandem wildtype Aβ) was a gift from Dr. Pedro Fernandez-Funez (University of Minnesota, USA)^21^. UAS-Mys (β1-integrin) was a gift from Dr. Rongwen Xi (NIBS, China)^86^. UAS-LanB1:RFP, UAS-Cg25C:RFP, UAS-LanB2:GFP, and UAS-secr:GFP (SP^Wg^.GFP) lines were a gift from Dr. José Carlos Pastor-Pareja (Tsinghua University, China)^25,87^. ElavGS driver line was a gift from Dr. Hervé Tricoire (CNRS, France)^88^. NsybGS driver line was a gift from Dr. Amita Sehgal (UPenn, USA)^89^. GliaGS driver line (GSG3285-1) was a gift from Dr. Minoru Saitoe (TMiMS, Japan) originally generated by Dr. Haig Keshishian (Yale, USA)^90^. Dilp2GS driver line (stock #3 on the second chromosome) was a gift from Dr. Heinrich Jasper (Buck Institute, USA)^91^.

Mutants and transgenic lines were backcrossed into the w^Dah^ *Wolbachia*-positive strain for at least six generations. Fly stocks were maintained and all experiments were conducted at 25°C on a 12h:12h light/dark circadian cycle at constant 65% humidity using standard sugar/yeast/agar (SYA) medium (containing 10% (w/v) brewer’s yeast, 5% (w/v) sucrose and 1.5% (w/v) agar)^92^. For all experiments involving Mifepristone, the steroid drug inducer ‘RU’ (RU486 – Sigma, Poole, Dorset, UK), the compound was dissolved in EtOH to make 100 mM stock solution and added to molten fly food for a final concentration of 200 μM. RU food was then dispensed into plastic vials, allowed to cool, then sealed and stored in a 4°C cold room. RU food was brought to room temperature before use with flies.

### Lifespan assay

Flies were reared at a standard density before being used for lifespan experiments as previously described^92,93^. All experiments were performed with flies (females and males) that were allowed 48 h to mate after emerging as adults. Flies were subsequently lightly anaesthetized with CO_2_, sorted into single sexes and counted at 15 per vial into 10 vials for a total of 150 flies per condition. In all cases, flies were transferred to fresh food at least three times a week, at which point deaths/censors were scored. For some survival assays, vials were kept in DrosoFlippers (drosoflipper.com) for ease of regular transfer to fresh vials. Microsoft Excel (lifespan template available at piperlab.org/resources/) was used to calculate survival proportions. Log-rank tests of survivorship curves were performed in Excel (Microsoft), and Cox proportional hazards analysis for multiple comparisons was performed in R Studio (R Core Team).

### Climbing assay

For each condition analysed in climbing assays, five vials of control food and five of food containing RU, each containing 15 flies, were housed side-by-side in a single DrosoFlipper. Flies were maintained as in lifespan studies and then were filmed for climbing assays once to twice per week. For climbing assays, the flies were kept in DrosoFlippers and transferred to empty vials on each side of the flipper, creating a standard vertical column 20cm in height for each set of flies. Flies were tapped to the bottom of the vials and allowed to climb upwards for 15 s before a still camera image was captured. The heights of individual flies were then assessed by manual multi-point selection in Fiji software^94^, with each height in pixels calibrated to a height in cm from a ruler placed next to the vials during filming.

### *Drosophila* rough eye phenotype analysis

Virgin females expressing Aβ^X2^ under the control of the GMR-Gal4 driver were crossed to males of the relevant genotype. Before imaging, flies were briefly snap frozen in liquid nitrogen to aid in correct positioning and avoid any movement artefacts. Eye images of 7-day-old flies were taken using a Leica M165 FC stereomicroscope equipped with a motorized stage and a multi-focus tool (Leica application suite software). For eye size analysis, wide-field images at the same magnification were taken of flies for each genotype and eye size was measured using Fiji^94^.

### Capillary feeder (CAFE) assay

A 7 mL bijou vial filled with 1 ml of (1%) agar, to ensure humid conditions, was sealed with Parafilm (Alpha Laboratories Ltd, Hampshire, UK). Four holes in the Parafilm that were equally spaced apart, were made using a 26-gauge needle to ensure adequate air circulation. Through the Parafilm was inserted a truncated 200 µl pipette tip which held a graduated 5 µl disposable glass capillary tube (Camag, Muttenz, Switzerland) containing liquid food (2% (w/v) yeast and 5% (w/v) sugar) supplemented with blue food dye (Langdale, Market Harborough, UK) to aid measurement of feeding. For all experiments, a mineral oil overlay (0.1 µl) was used to minimize evaporation. Food ingestion was measured every 24-hr. Each experiment included an identical, CAFE chamber without flies to determine evaporative losses (typically 10% of ingested volumes), which were subtracted from experimental readings^60^.

### Brain dissection, immunohistochemistry, and imaging of the fly brain

We followed the dissection and staining protocol for Aβ detection in the adult *Drosophila* brain from Ray *et al*. 2017^18^. Using forceps, fly heads were removed from the bodies in cold PBS containing 4% paraformaldehyde (Pierce, Thermo Fisher) + 0.01% Triton X-100 (Sigma). The proboscis was then removed from the fly head to allow fixative and detergent to fix and permeabilise the fly brain. The fly heads were transferred to a 0.5 mL microcentrifuge tube, fixed in 400 μL 4% paraformaldehyde + 0.01% Triton X-100 and rocked for 16 mins at room temperature. The heads were washed with PBST-1 (PBS containing 0.01% Triton-X-100) for 3 x 2 mins at room temperature. The brains were dissected from the fly heads in ice cold PBST-1 in a dissection dish under a light stereomicroscope. The brains were transferred to a new 0.5 mL microcentrifuge tube and fixed in 400 μL 4% paraformaldehyde + 0.1% Triton X-100 and rocked for 20 mins at room temperature. The brains were then washed in PBST-2 (PBS containing 0.1% Triton X-100) for 3 x 2 mins and blocked with SeaBlock blocking buffer (Thermo Fisher) for 15 mins at room temperature. SeaBlock blocking buffer was removed and the brains were incubated in primary antibody in PBST-2 overnight, washed in PBST-2 for 3 x 20 mins and incubated in secondary antibody in PBST-2 for 1 hour in the dark. The brains were washed in PBST-2 for 3 x 20 mins at room temperature in the dark and mounted on a microscope slide using Vectashield mounting media containing DAPI (Vector Labs). The following antibodies were used. Primary antibodies: mouse anti-Aβ 6E10 (1:500; BioLegend #803001). Secondary antibodies: Alexa Fluor 488 donkey anti-mouse (1:400; Thermo Fisher #A21202), Alexa Fluor 594 goat anti-mouse (1:400; Thermo Fisher #A11005). Images were captured with a Zeiss LSM 700 confocal laser scanning microscope (Zeiss, Germany) or with a Leica TCS8 confocal microscope (Leica, Wetzlar, Germany) with a 20x or 63x oil immersion objective. Images were taken as stacks and are shown as single sections or maximum intensity projections of the complete stack. All images for one experiment were taken at the same settings.

### Live 2-photon imaging

Adult flies were fixed ventral side down to microscope slides using dental composite (3M Espe Sinfony Enamel Effect Material) and cured using a curing light (3TECH LED-1007 #104-0028). A small piece of cuticle was removed from the posterior side of the head (cuticular window) to reveal the fly brain. GFP fluorescence resulting from LanB2:GFP expression in Dilp2 neurons was imaged with a Leica TCS SP8 MP 2-photon microscope (Leica, Wetzlar, Germany).

### Quantitative real-time PCR (qPCR)

Total RNA was isolated from adult fly heads using standard TRIzol (Invitrogen) protocols. RNA samples were treated with Turbo DNAse (Invitrogen) and converted to cDNA using oligo-dT primers and Superscript II reverse transcriptase (Invitrogen). Quantitative RT– PCR was performed using Power SYBR Green PCR Master Mix (ABI) in the Quant Studio 6 Flex system (Applied Biosystems, Thermo Fisher Scientific). Each sample was analyzed in duplicate, and values are the mean of four or five independent biological repeats. Relative quantities of transcripts were determined using the relative standard curve method normalized to Tub84B or Rp49. The following primer sequences (Eurofins, UK) were used in the analysis: Aβ_42_: 5’-CGATCCTTCTCCTGCTAACC-3’, 5’- CACCATCAAGCCAATAATCG-3’; LanB1 (PP29286): 5’-CTCGCCGGAGAGATTCTGC-3’, 5’-TTGTACGGATCATGCTTGGTC-3’; cdc14 (PP30015): 5’- GACTTTGGTCCGCTCAACATA-3’, 5’-CGGATTCATGGAGGTGTAGTGA-3’; BiP: 5’- TCTTGTACACACCAACGCAGG-3’, 5’-CAAGGAGCTGGGCACAGTGA-3’; Tub84B: 5’- TGGGCCCGTCTGGACCACAA-3’, 5’-TCGCCGTCACCGGAGTCCAT-3’; Rp49: 5’- GACAATCTCCTTGCGCTTCT-3’, 5’-CCAGTCGGATCGATATGCTAA-3’. LanB1 (PP29286) and cdc14 (PP30015) primer sequences were obtained from the Fly Primer Bank^95^ (flyrnai.org/flyprimerbank).

### Quantification of total, soluble and aggregated Aβ42

To extract total Aβ_42_, five fly heads were homogenised in 50 μl GnHCl extraction buffer (5 M Guanidinium HCl, 50 mM HEPES pH 7.3, protease inhibitor cocktail (Sigma, P8340) and 5 mM EDTA), centrifuged at 21,000×g for 5 min at 4°C, and cleared supernatant retained as the total fly Aβ_42_ sample. Alternatively, soluble and insoluble pools of Aβ_42_ were extracted using a protocol adapted from previously published methods^56,96^: 50 fly heads were homogenised in 50 μL tissue homogenisation buffer (250 mM sucrose, 20 mM Tris base, 1 mM EDTA, 1 mM EGTA, protease inhibitor cocktail (Sigma)) then mixed further with 200 μl DEA buffer (0.4% DEA, 100 mM NaCl and protease inhibitor cocktail). Samples were centrifuged at 135,000 × g for 1 hr at 4°C (Beckman OptimaTM Max centrifuge, TLA120.1 rotor), and supernatant retained as the cytosolic, soluble Aβ_42_ fraction. Pellets were further resuspended in 400 μl ice-cold formic acid (70%) and sonicated for 2× 30 s on ice. Samples were re-centrifuged at 135,000 × g for 1 hr at 4°C, then 210 μl of supernatant diluted with 4 ml FA neutralisation buffer (1 M Tris base, 0.5 M Na2 HPO4, 0.05% NaN3) and retained as the aggregated, formic acid-extractable Aβ_42_ fraction. Total, soluble or aggregated Aβ_42_ content was measured using the Ultrasensitive hAmyloid-β_42_ ELISA kit (Thermo Fisher, # KHB3544). Total and soluble Aβ_42_ samples were diluted 1:10, and aggregated Aβ_42_ samples diluted 1:5 in sample/standard dilution buffer and the ELISA performed according to the manufacturers’ instructions. Protein extracts were quantified using the BCA Protein Assay Kit (Pierce) and the amount of Aβ_42_ in each sample expressed as a ratio of the total protein content (pmol/g total protein).

### Western blots

For protein extracts, fly heads (10-20 per biological sample) were homogenised in 1x NuPAGE LDS sample buffer (Thermo Fisher) with 200 mM DTT (Sigma) and boiled at 95°C for 5 min. With the assumption that fly heads of the same sex contain comparable levels of total protein, we lysed 1 fly head per 10 μL sample buffer. Equal quantities of protein for each sample were then separated on 4%–12% NuPAGE Bis-Tris gels (Invitrogen) and transferred to a nitrocellulose membrane (GE Healthcare). For Aβ westerns antigen retrieval was necessary, so membranes were boiled in 1X PBS for 5 min in a microwave. Membranes were blocked in 5% bovine serum albumin (BSA, Sigma) in Tris-buffered saline (TBS) with 0.1% Tween-20 (TBST) for 1 hr at room temperature, after which they were probed with primary antibodies overnight at 4°C. The following primary antibodies were used: mouse anti-Aβ 6E10 (1:500; BioLegend #803001 previously #39320), rabbit anti-BiP (1:1000; Novus Biologicals #NBP1-06274), rabbit anti-Actin (1:5,000; Abcam #ab1801), mouse anti-Tubulin (1:2000, Sigma #T6199). Membranes were then probed with secondary anti-mouse HRP (1:10,000; Abcam #ab6789) or anti-rabbit HRP antibodies (1:10,000; Abcam #ab6721) for 1 hr at room temperature. Blots were developed using Luminata Crescendo (Millipore) and the ImageQuant LAS 4000 system. Densitometric analysis of blot images was carried out using Fiji software^94^.

### Single-cell transcriptomic data for the adult fly brain

Single-cell atlas images were obtained by gene name searches in *SCope*^59^, the vizualisation and analysis tool from the Aerts lab (http://scope.aertslab.org/).

### Mice

P10 wildtype mice (C57/BL6J) were used for slice culture experiments. Animals were culled by humane schedule one methods and brain tissue removed for the generation of organotypic slice cultures. All animal experiments conformed to national and institutional guidelines including the Animals [Scientific Procedures Act] 1986 (UK), and the Council Directive 2010/63EU of the European Parliament and the Council of 22 September 2010 on the protection of animals used for scientific purposes, and had full Home Office ethical approval. Mice were bred in-house and group-housed on a 12h/12h light/dark cycle with *ad libitum* access to food and water.

### Organotypic slice cultures

Organotypic cultures of the hippocampus and surrounding cortex were taken from humanely sacrificed P10 mouse pups of either sex according to previously described protocols^97–99^. Briefly, brains were rapidly removed and kept in dissection buffer (Earle’s Balanced Salt Solution (EBSS) + 25 mM HEPES + 1X Penicillin/Streptomycin) on ice. From this point, until plating, all equipment and tissue was kept ice cold. Brains were bisected at the midline then the cut sides glued (Loctite super glue), face down onto a vibratome stage and flooded with dissection media. 350 μm sagittal slices (6 per brain) were taken using a Leica VT1200S Vibratome; the hippocampus with surrounding cortex was dissected out using sterile syringe needles. The dissected slices were then transferred (using a sterile 3 ml plastic pipette - modified to widen the opening) to Falcon tubes full of ice-cold dissection medium and stored until plating. To plate, slices were transferred (3 slices from the same pup per dish, 2 dishes per pup (split randomly between control/ test conditions)) onto sterile 0.4 μm pore membranes (Millipore #PICM0RG50) in 35 mm culture dishes (Nunc). Inserts were kept in 1 ml of maintenance medium (50% MEM with Glutamax-1 (Life Tech:42360-024), 25% Heat-inactivated horse serum (Life Tech: 26050-070), 23% EBSS (Life Tech: 24010-043), 0.65% D-Glucose (Sigma: G8270), 2% Penicillin-Streptomycin (Life Tech: 15140-122) and 6 units/ml Nystatin (Sigma: N1638)) and cultures were maintained in incubators at 37°C, 5% CO2 for 4 weeks. Two 100% medium exchanges occurred (5 hr after plating and 4 days *in vitro* and a 50% media exchange occurred each week thereafter. At 14 days *in vitro*, LamB1 or control lentivirus was applied at a viral titer of 8.33×10^7^ per dish (diluted in the culture medium with a few drops added on top of the slice to ensure compete perfusion of the tissue). Culture medium was changed 24 hours later, and cultures maintained for a further 14 days *in vitro* before fixation.

### Immunofluorescence staining of organotypics

Slice cultures were fixed for 20 mins in 4% paraformaldehyde in PBS. Slices were washed twice in PBS, blocked for 1 h in blocking solution (PBS with 0.5% Triton X-100 and 3% Goat Serum) then incubated in 200 μl primary antibody diluted in blocking solution overnight at 4°C with shaking. Slices were washed 3 times in PBS before being incubated (2 hr, RT in the dark) with secondary antibodies in blocking solution. In the second of the final 3 PBS washes, slices were incubated with DAPI (Sigma, 1:10,000 in PBS). Images were captured using a Leica TCS8 confocal microscope (Leica, Wetzlar, Germany). Primary antibodies used: rat Lamb1 (1:250; Abcam #ab44941), rabbit Calnexin (1:250; Abcam #ab13504). Secondary antibodies used: goat anti-rabbit Alexa Fluor 594 (1:400; Invitrogen #A11037) and goat anti-rat Alexa Fluor 647 (1:400; Invitrogen #A21247).

### Statistics

Data were grouped for each genotype and the mean (+/- SEM) calculated. Log-rank, Cox Proportional Hazard, Analysis of Variances (ANOVA) and Tukey’s HSD (honestly significant difference) post-hoc analyses were performed. Statistical analyses were performed in Excel (Microsoft) or Prism (GraphPad, La Jolla, CA), except for Cox Proportional Hazards which were performed in R Studio (R Core Team). A statistical difference of p<0.05 was regarded as significant.

## Acknowledgements

We thank Dr Elizabeth Catterson, Dr Teresa Niccoli, and Dr John Labbadia for helpful comments, and Dr Adam Dobson for statistical help. We are grateful to past and present members of the Partridge and Spires-Jones labs for helpful discussions. We also thank Giovanna Vinti for maintenance of the *Drosophila* laboratory facilities and Michael Wright for administrative support in the Institute of Healthy Ageing at UCL. We also thank the wider fly community for generous sharing of reagents and stocks, particularly the Bloomington *Drosophila* Stock Center (NIH P40OD018537) and the Vienna *Drosophila* Resource Center. This work was funded by a Wellcome Trust Strategic Award (098565) and an ERC Award (ALZSYN 681181).

## Author contributions

Conceived and designed the experiments: JHC LP. Performed the experiments: JC LM SA SJM SM MCD AR NSW MLA MA CD. Analysed the data: JHC. Contributed reagents/materials/analysis tools: JHC TLSJ LP. Wrote the paper: JHC LP.

## Supplementary Figures

**Supplementary Fig. 1.**
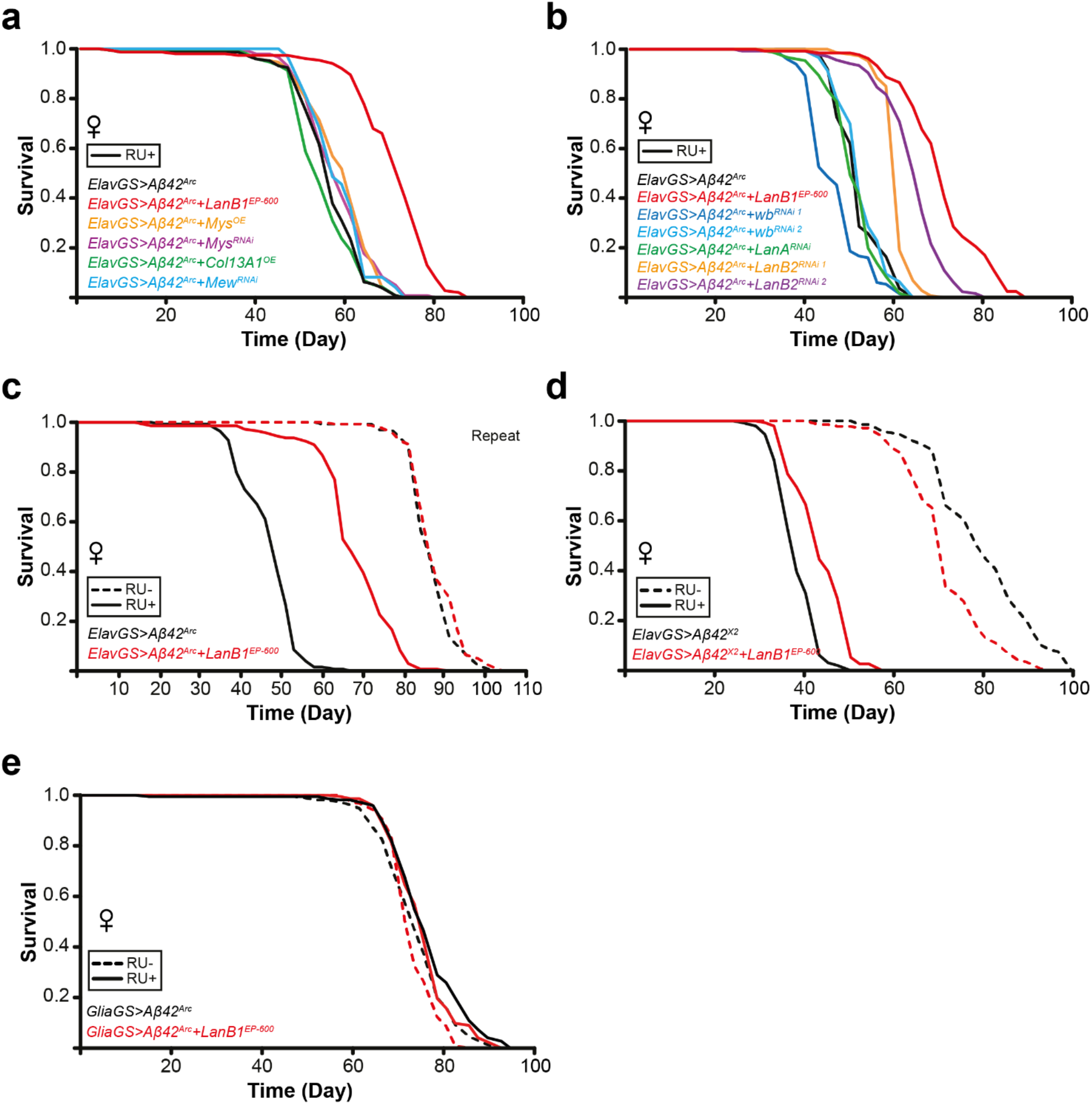
Uncovering a role for LanB1 in the amelioration of Aβ toxicity. **a** ECM-related transgenic lines were crossed with flies expressing pan-neuronal Aβ^Arc^ with the ElavGS driver. LanB1 co-expression significantly rescued the short-lifespan phenotype compared to the induced Aβ^Arc^-alone control (p = 2.86 x 10^−44^; log rank test). **b** Additional Laminin transgenic lines were examined for their effect on Aβ toxicity. LanB1 co-expression significantly rescued toxicity compared to induced controls (p = 1.08 x 10^−54^; log rank test). Both independent LanB2 RNAi lines also exhibited significant rescue of Aβ toxicity (LanB2^RNAi 1^ p = 7.94 x 10^−35^; LanB2^RNAi 2^ p = 4.13 x 10^−41^ log rank test). **c** Induction of Aβ^Arc^ significantly (p = 8.34 x 10^−67^; log rank test) shortened lifespan compared to uninduced controls. LanB1 and Aβ^Arc^ co-expression resulted in a significant rescue (p = 1.48 x 10^−56^; log rank test) of the short-lived phenotype. **d** Expression of two copies of wildtype Aβ_42_ (Aβ^X2^) in neurons. Aβ^X2^ significantly (p = 4.52 x 10^−68^; log rank test) shortened lifespan compared to uninduced controls. There was a significant difference between uninduced control lines (p = 4.00 x 10^−12^; log rank test), but in the reverse direction to the rescue. LanB1 and Aβ^X2^ co-expression resulted in a significant rescue (p = 6.47 x 10^−15^; log rank test). **e** Glial expression of Aβ was not toxic. Using a glial GeneSwitch driver (GliaGS), flies with adult-onset glial Aβ expression had slightly extended lifespan compared to uninduced controls (Aβ p = 0.0074; Aβ+LanB1 p = 0.0074; log rank test). LanB1 co-expression did not affect this extension (comparison in induced conditions p = 0.14; log rank test). For lifespan experiments n = 150 flies per condition.

**Supplementary Fig. 2.**
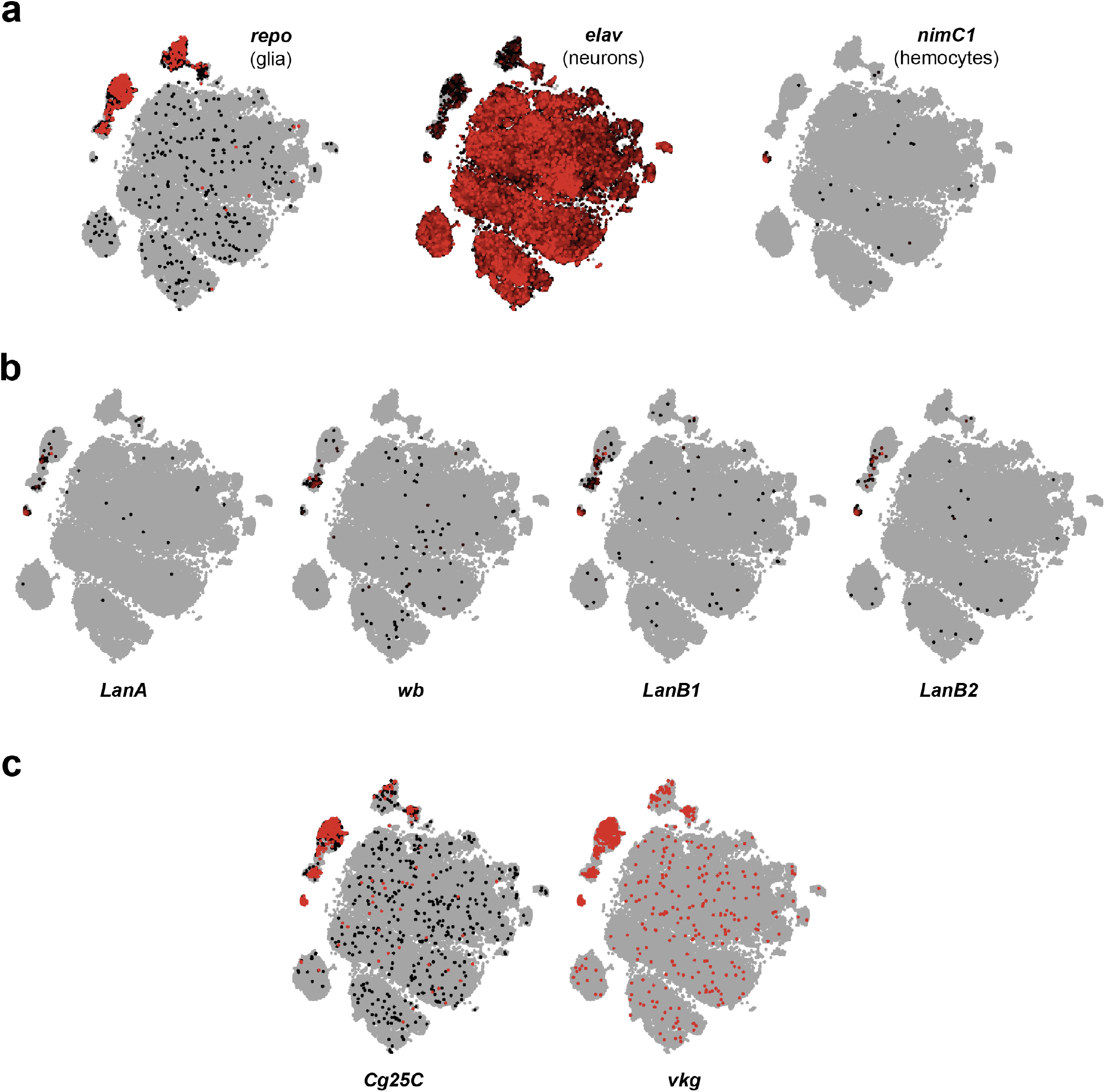
Single-cell transcriptome atlas of the adult *Drosophila* brain shows that laminin- and collagen IV-subunit expression is restricted to haemocytes and a subset of glial cells. **a** Cell types in the brain can be distinguished by specific gene expression. Glial cells are specified by *repo*, neurons by *elav*, and haemocytes by *nimC1* expression. **b** All four laminin subunits in *Drosophila* (LanA, wb, LanB1, LanB2) are expressed in haemocytes and a subset of glial cells but are not substantially expressed in neurons. **c** The two collagen IV subunits (Cg25C, vkg) are highly expressed in haemocytes and glial cells with some neuronal expression. Atlas images were obtained by gene name searches in *SCope*, the vizualisation and analysis tool from the Aerts lab (http://scope.aertslab.org/).

**Supplementary Fig. 3.**
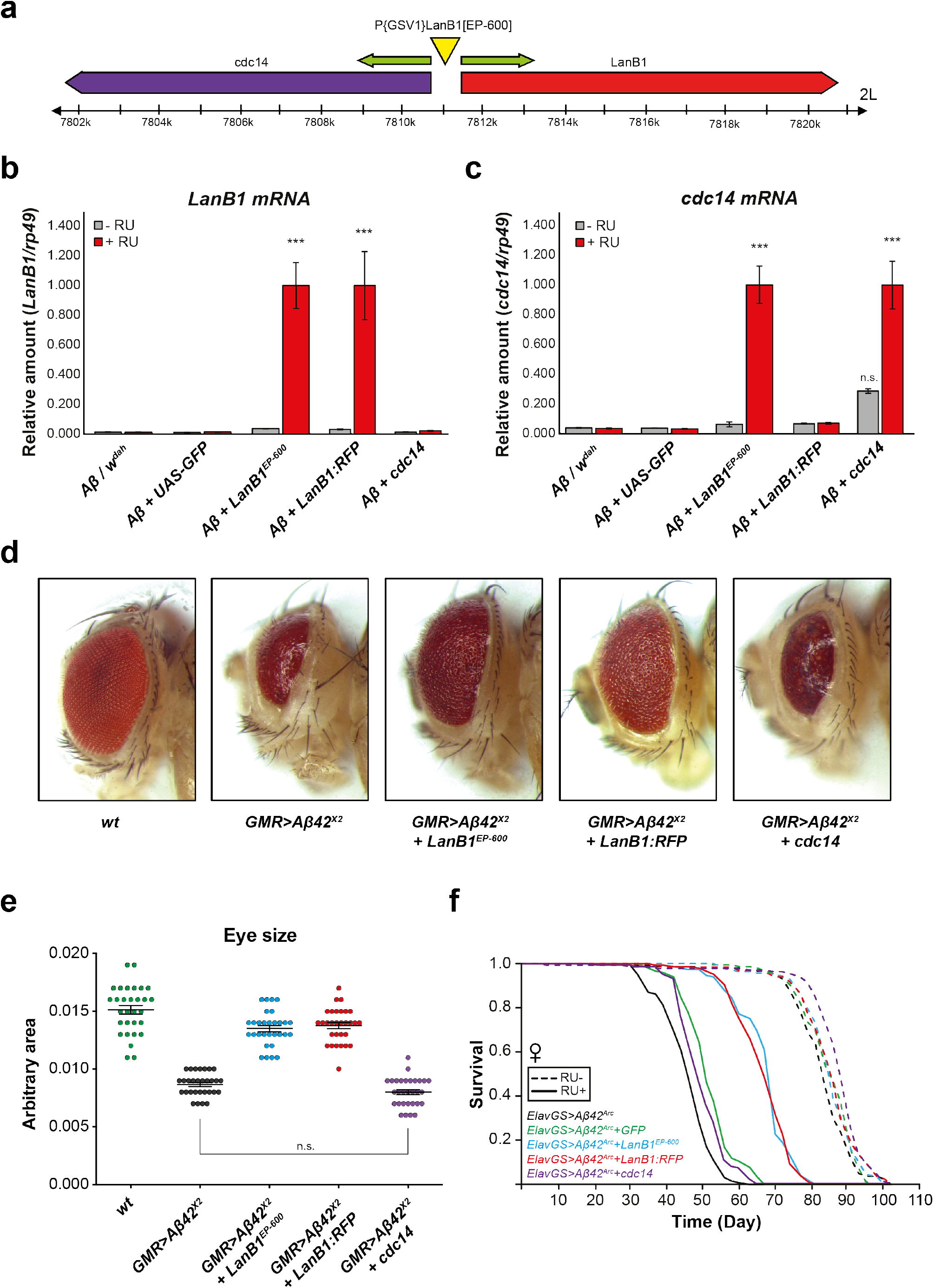
Expression of *LanB1* and not *cdc14*, an adjacent gene in the opposite orientation, rescued Aβ toxicity. **a** Schematic of the LanB1^EP-600^ locus located 548 bases upstream of the 5’ UTR of the LanB1 gene. The P{GSV1} element contains UAS sequences at both ends oriented outwards. **b** *LanB1* was significantly upregulated in the heads of UAS-LanB1^EP-600^ and UAS-LanB1:RFP flies compared to uninduced controls (p < 0.0001; one-way ANOVA with Tukey’s post-hoc test). **c** *Cdc14* was significantly up-regulated in the heads of UAS-LanB1^EP-600^ and in UAS-cdc14 (cdc14^EY10303^) heads compared to uninduced controls (p < 0.0001; one-way ANOVA with Tukey’s post-hoc test). Cdc14 up-regulation was not observed in UAS-LanB1:RFP heads. Mild up-regulation in uninduced UAS-cdc14 heads may be due to compensatory up-regulation as the cdc14^EY10303^ P-element is inserted 395 bases into the 5’ UTR of cdc14. Relative qPCR values were normalized to *rp49* expression. Data are shown as mean ± SEM (n = 4 biological replicates per condition). **d** Co-expression of Aβ^X2^ with LanB1 (via UAS-LanB1^EP-600^ and UAS-LanB1:RFP) using the eye-specific GMR-GAL4 driver rescued most of the size and organisation of the eye. Cdc14 did not rescue Aβ^X2^ toxicity. For display purposes, three micrographs in **d** are the same as those in Fig 1a. **e** Quantification of eye sizes in **d**. There was no significant difference in eye size between Aβ^X2^ alone or with cdc14 co-expression. LanB1 (via UAS-LanB1^EP-600^ and UAS-LanB1:RFP) significantly rescued eye size (p < 0.0001; one-way ANOVA with Tukey’s post-hoc test). Data are shown as mean ± SEM (n = 30 eyes measured per condition). **f** The considerable rescue from Aβ toxicity by LanB1^EP-600^ was due to LanB1 and not cdc14. Although co-expression of cytoplasmic GFP (BDSC #1521) or cdc14 with Aβ^Arc^ led to small but significant rescues of lifespan compared to Aβ^Arc^ alone controls (cytoplasmic GFP, p = 2.44 x 10^−12^; cdc14, p = 2.64 x 10^−6^ log rank test), LanB1 and Aβ^Arc^ co-expression (via UAS-LanB1^EP-600^ and UAS-LanB1:RFP) exhibited equivalently large rescues of Aβ toxicity (LanB1^EP-600^, p = 1.39 x 10^− 61^; LanB1:RFP, p = 2.85 x 10^−62^ log rank test). n = 150 flies per condition.

**Supplementary Fig. 4.**
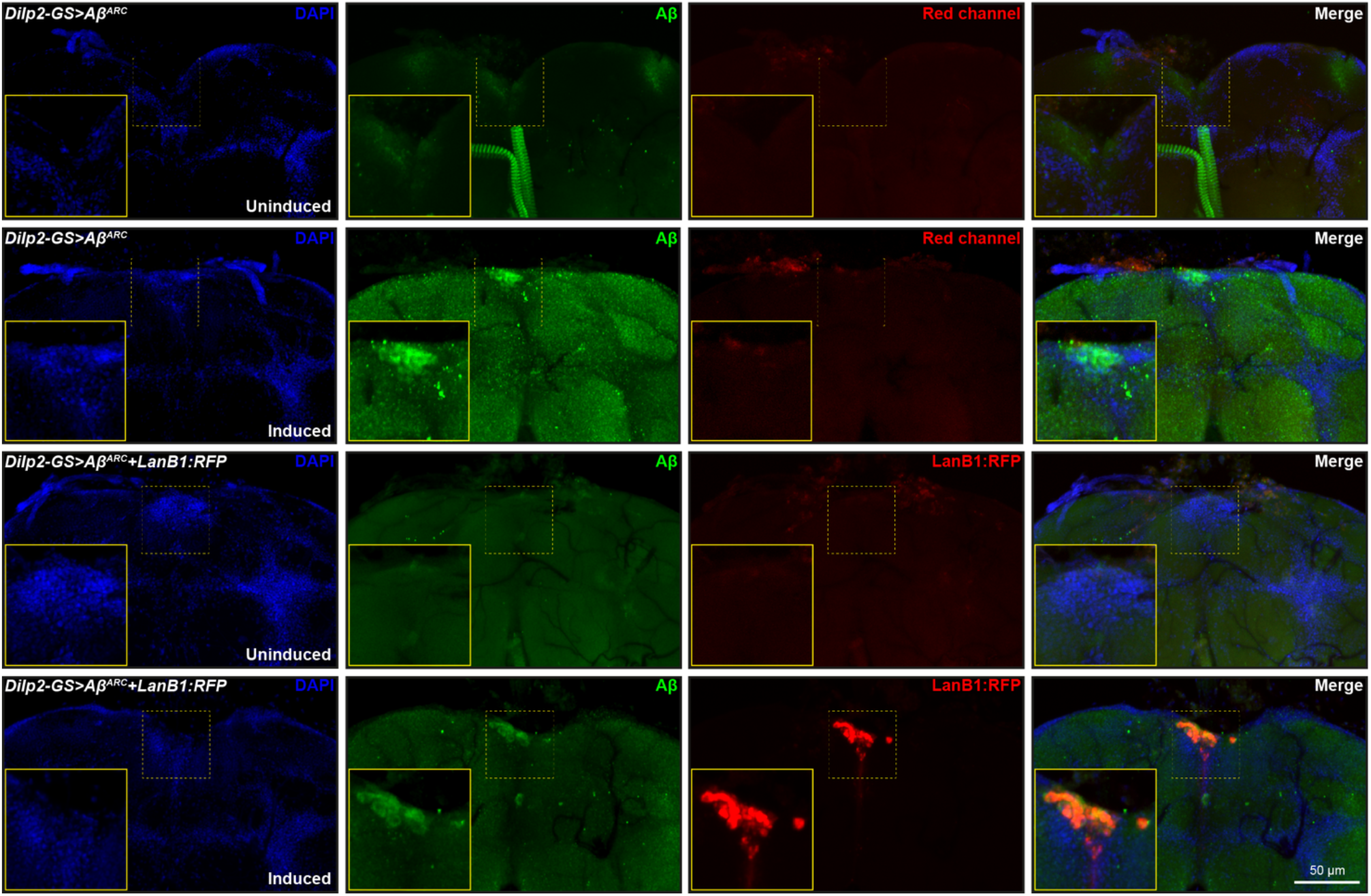
Inducible expression of Aβ and LanB1 in Dilp2 neurons using the Dilp2-GS driver. Without RU induction, there was no induction of Aβ and/or LanB1 expression. When fed RU, Aβ and LanB1 were expressed. Representative confocal fluorescence z projections taken at 20x magnification of whole brains from 21-day-old female flies stained with Aβ (6E10 – green) and DAPI (blue). Yellow box inset shows a higher magnification of the Dilp2 neuron cluster area in the dorsal brain. Endogenous fluorescence (i.e. without staining) of LanB1:RFP is shown. Genotypes: *Dilp2-GS>Aβ^Arc^*; and *Dilp2-GS>Aβ^Arc^ + LanB1:RFP*. Scale bar, 50 μm.

**Supplementary Fig. 5.**
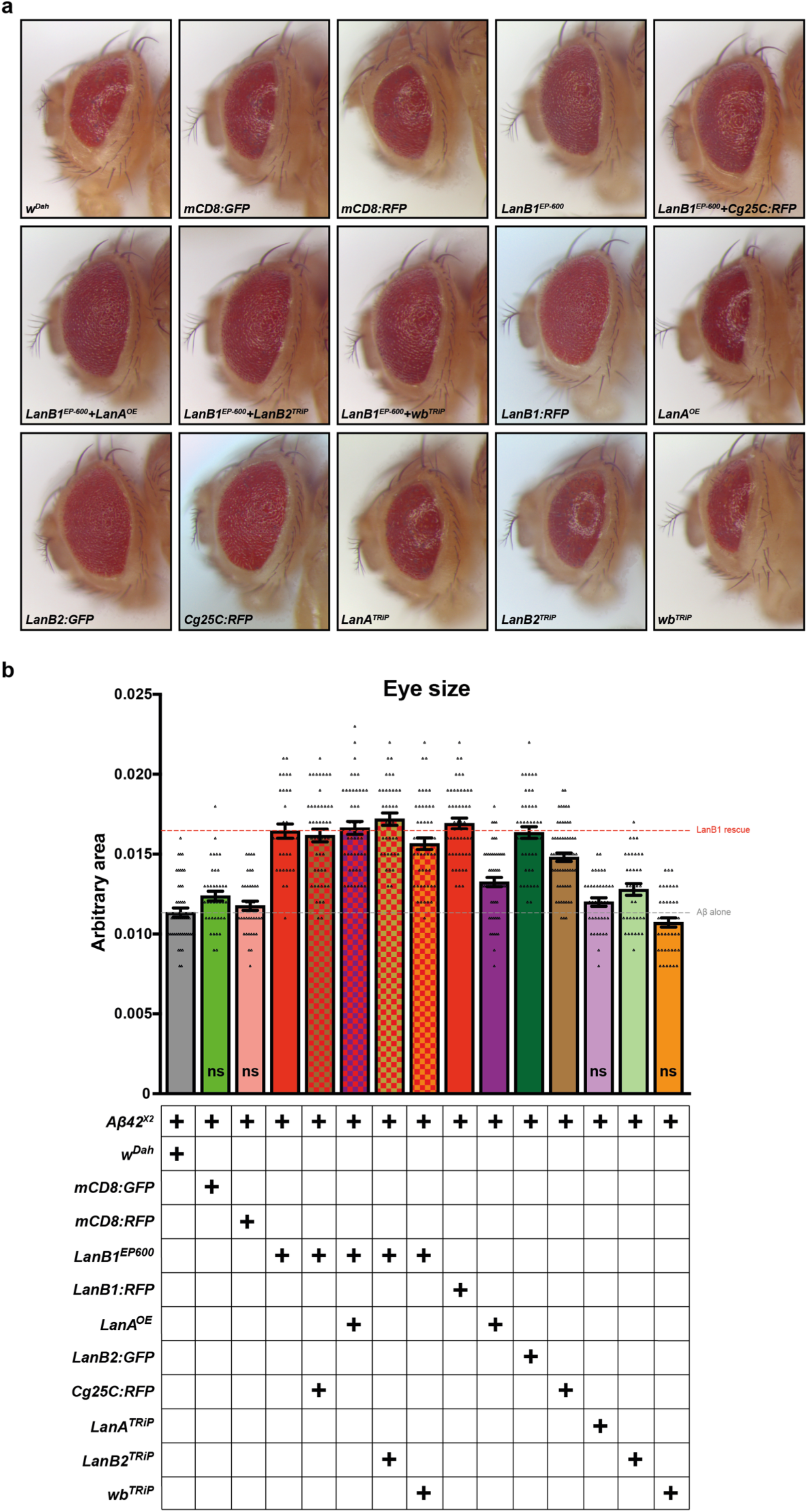
Laminin-induced rescue of Aβ toxicity in the developing eye is not affected by modulation of other laminin subunits. **a** All flies shown expressed Aβ^X2^ using GMR-GAL4. The degree of rescue from LanB1 over-expression (via UAS-LanB1^EP-600^ and UAS-LanB1:RFP) was not affected by expression of the indicated genes. **b** Quantification of eye sizes in **a**. Eye size for each condition was quantified and compared to the *ElavGS>Aβ^X2^/w^Dah^* control. Over-expression or knockdown of other Laminin subunits in combination with LanB1 had no significant effect on the rescue. Conditions marked ‘ns’ were not significantly different compared to controls, while the other lines were significantly different (one-way ANOVA with Tukey’s post-hoc test). Plus (**+**) symbol indicates presence of labelled transgene. Bars containing red indicate presence of a LanB1 over-expression line. Dashed grey line indicates the Aβ control, while dashed red line indicates the LanB1-rescue level. ‘TRiP’ indicates an RNAi transgene. Data are shown as mean ± SEM (n = 34-62 eyes measured per condition).

**Supplementary Fig. 6.**
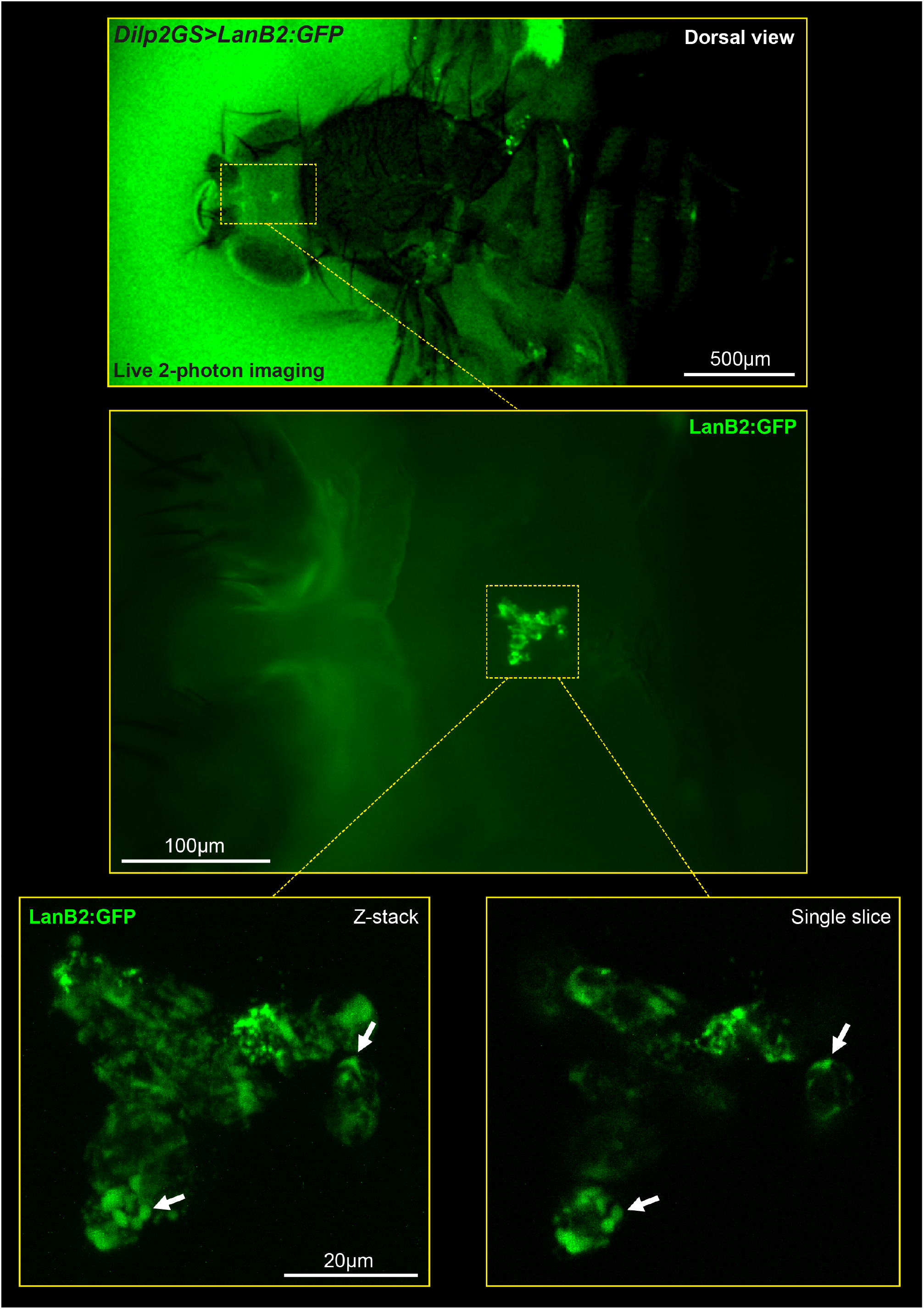
Live 2-photon imaging of a fly producing LanB2:GFP in Dilp2 neurons. LanB2 accumulated intracellularly in discrete compartments. *Top*, dorsal view of fly with cuticle removed to visualize the brain. To prevent dehydration, saline gel was added with a small coverslip on top. *Middle*, closer dorsal view of live fly brain. *Bottom left*, z-projection of the cell bodies of the Dilp2 neurons expressing LanB2:GFP. *Bottom right*, single slice of these cell bodies showing LanB2 accumulated in discrete cellular compartments and not diffusely in the cytoplasm. Arrows show discrete intracellular accumulations of LanB2:GFP. Genotype: *Dilp2-GS>LanB2:GFP*. In all images anterior is left, posterior is right. Scale bar, top 500 μm, middle 100 μm, bottom, 20 μm.

**Supplementary Fig. 7.**
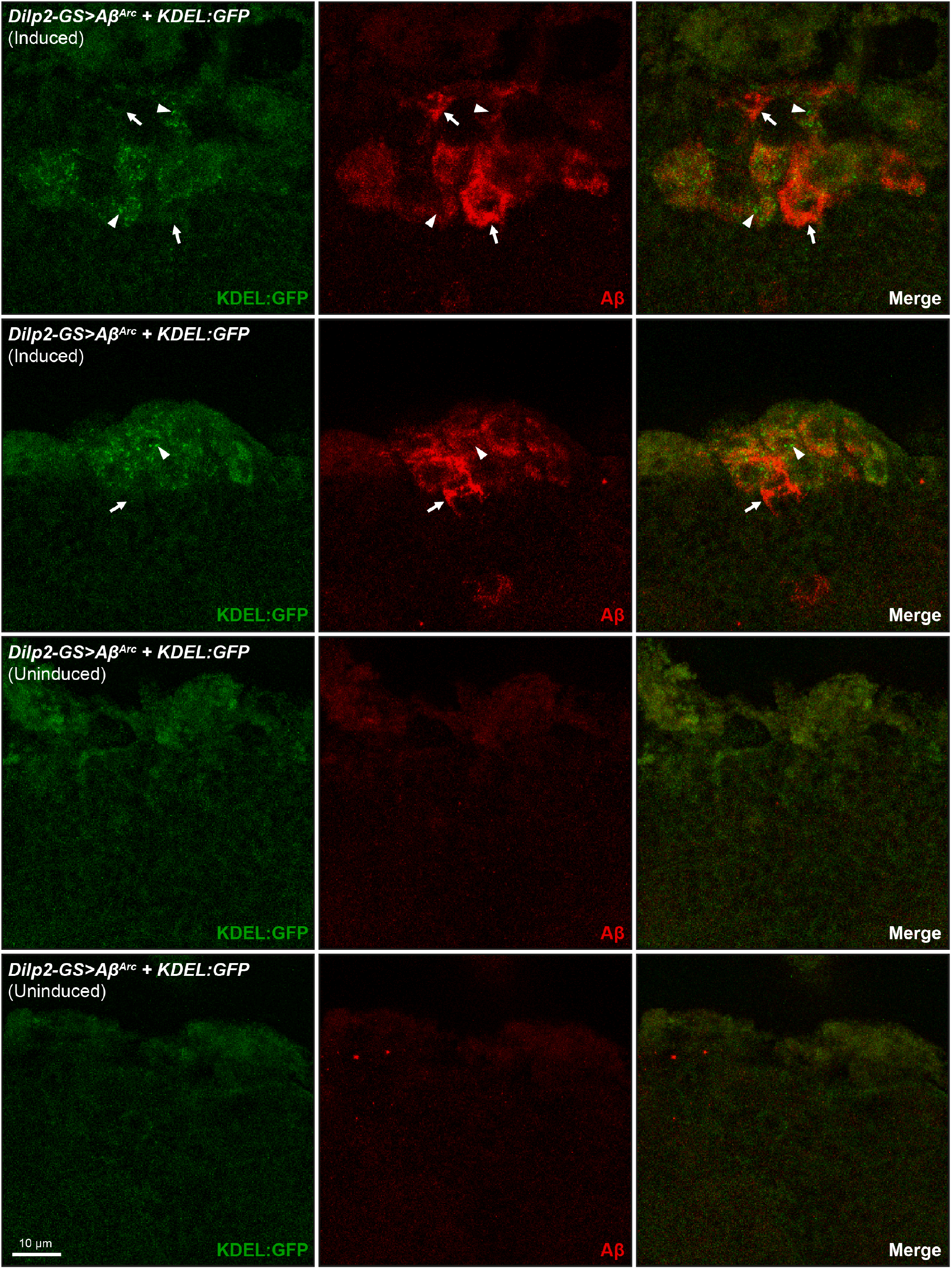
Aβ does not accumulate in the ER. Inducible expression of Aβ and the ER marker, KDEL:GFP, in Dilp2 neurons using the Dilp2-GS driver. *Top two rows*, when fed RU, Aβ and KDEL:GFP were present in the cell bodies of Dilp2 neurons. Arrows highlight areas with no overlap of Aβ and KDEL:GFP. Arrowheads highlight punctate areas of ER with no corresponding Aβ expression. *Bottom two rows*, without RU486 induction, there was no induction of Aβ and/or KDEL:GFP expression. Representative confocal fluorescence z projections taken at 63x magnification from 21-day-old female flies stained with Aβ (6E10 – green). Endogenous fluorescence (i.e. without staining) of KDEL:GFP is shown. Genotype: *Dilp2-GS>Aβ^Arc^ + KDEL:GFP*. Scale bar, 10 μm.

**Supplementary Fig. 8.**
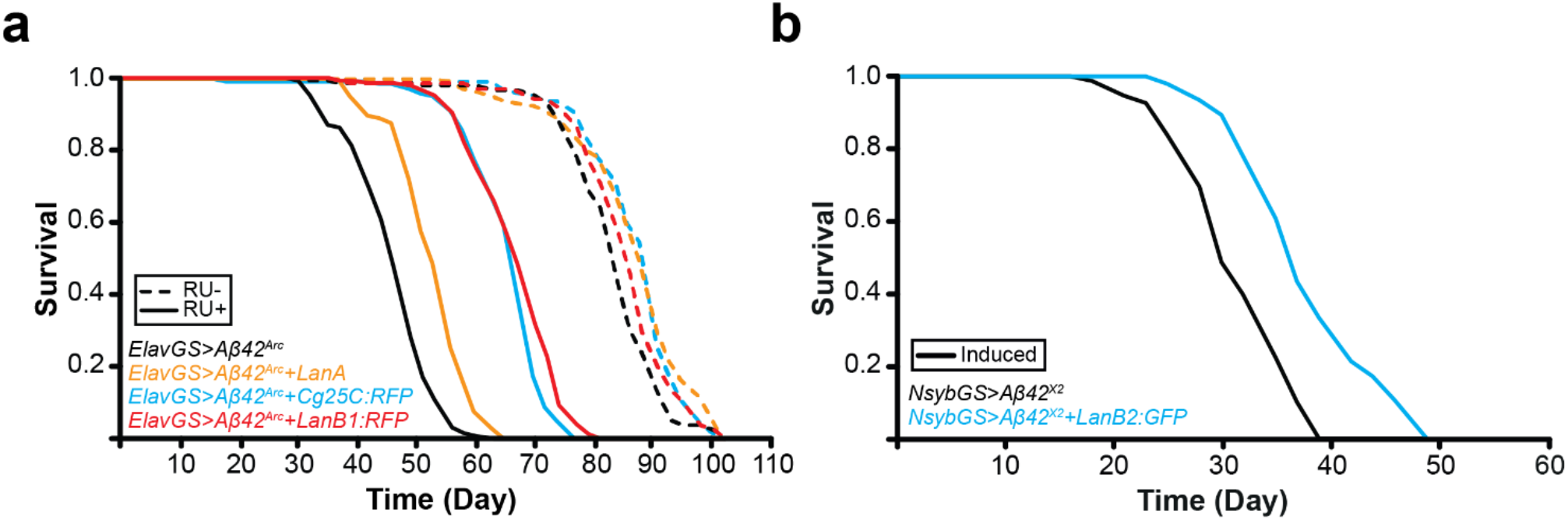
Laminin γ-chain (LanB2) also rescued Aβ toxicity. **a** LanB1, Cg25C, and LanA rescued Aβ toxicity (LanB1, p = 2.85 x 10^−62^; Cg25C, p = 3.43 x 10^−63^; LanA, p = 1.58 x 10^−17^; log rank vs *ElavGS>Aβ^Arc^* alone). There were significant extensions of lifespan in the uninduced controls (LanB1, p = 0.036; Cg25C, p = 7.50 x 10^−06^; LanA, p = 7.33 x 10^−06^; log rank vs *ElavGS>Aβ^Arc^* alone). **b** LanB2 and Aβ^X2^ co-expression using NsybGS resulted in a significant rescue (p = 1.00 x 10^−20^; log rank test) compared to induced Aβ^X2^ controls. For all lifespan experiments n = 150 flies per condition.

